# Loss of neuropeptidergic regulation of cholinergic transmission induces CaV1-mediated homeostatic compensation in muscle cells

**DOI:** 10.1101/2024.09.10.612183

**Authors:** Jiajie Shao, Jana F. Liewald, Wagner Steuer Costa, Christiane Ruse, Jens Gruber, Mohammad Suleiman Djamshedzad, Wulf Gebhardt, Alexander Gottschalk

**Affiliations:** Institute of Biophysical Chemistry, Faculty of Molecular Sciences, Goethe University, Max-von-Laue-Strasse 9, D-60438 Frankfurt, Germany; Buchmann Institute for Molecular Life Sciences, Goethe University, Max-von-Laue-Strasse 15, D-60438 Frankfurt, Germany

## Abstract

Chemical synaptic transmission at the neuromuscular junction (NMJ) is regulated by electrical activity of the motor circuit, but may also be affected by neuromodulation. Here, we assess the role of neuropeptide signaling in the plasticity of NMJ function in *Caenorhabditis elegans*. We show that the CAPS (Ca^2+^-dependent activator protein for secretion) ortholog UNC-31, which regulates the exocytosis of dense core vesicles (DCVs), affects both pre- and post-synaptic functional properties, as well as NMJ-mediated locomotion. Despite reduced evoked acetylcholine transmission, the loss of *unc-31* results in a more vigorous response to presynaptic stimulation, i.e., enhanced muscle contraction and Ca^2+^ transients. Based on expression profiles, we identified neuropeptides involved in both cholinergic (FLP-6, NLP-9, NLP-21 and NLP-38) and GABAergic motor neurons (FLP-15, NLP-15), that mediate normal transmission at the NMJ. In the absence of these peptides, neurons fail to upregulate their transmitter output in response to increased cAMP signaling. We also identified proprotein convertases encoded by *aex-5/kpc-3* and *egl-3/kpc-2* that act synergistically to generate these neuropeptides. We propose that postsynaptic homeostatic scaling, mediated by increased muscle excitability, could compensate for the reduced cholinergic transmission in mutants affected for neuropeptide signaling, thus maintaining net synaptic strength. We show that in the absence of UNC-31 muscle excitability is modulated by upregulating the expression of the muscular L-type voltage gated Ca^2+^ channel EGL-19 (CaV1). Collectively, our results unveil a role for neuropeptidergic regulation in synaptic plasticity, linking changes in presynaptic transmission to compensatory changes in muscle excitability.

## INTRODUCTION

Chemical synaptic transmission at neuromuscular junctions is orchestrated by networks of motor neurons (MNs) and pre-motor interneurons, and involves pattern generation such that coordinated movement and also locomotion can occur (Marder and Bucher, 2001; Zhen and Samuel, 2015). The patterns of electrical activity of the MNs lead to the fusion of synaptic vesicles (SVs) and release of neurotransmitter (Sudhof, 2013). The rate of SV fusion depends on the extent and duration of depolarization of the synaptic terminal, which triggers voltage-gated Ca^2+^ channels (VGCCs) (Nanou and Catterall, 2018). It also depends on the mobilization of SVs from the reserve pool, i.e. when more SVs are mobilized, depolarization can cause the fusion of SVs at a higher rate. These mechanisms are subject to plasticity, regulating synaptic transmission (Zucker and Regehr, 2002; Davis, 2013; Kavalali, 2015; Nanou and Catterall, 2018; Reshetniak and Rizzoli, 2021; Silva et al., 2021; Zhang et al., 2022), and also enable presynaptic homeostatic upscaling. This can occur in situations where output of transmitter is affected, or in response to (reduced or missing) retrograde signaling from the postsynaptic side, when postsynaptic neurotransmitter detection is compromised. This is well characterized for the neuromuscular junction of *Drosophila* (Orr et al., 2017a; Orr et al., 2017b; Orr et al., 2022). There are also mechanisms that involve postsynaptic homeostatic scaling in response to a less than normal level of presynaptic transmitter release. This was shown in cultured neurons when presynaptic firing was chronically blocked using tetrodotoxin, or increased by GABA blockers, and caused an increase or decrease in miniature postsynaptic current amplitude, respectively (Turrigiano et al., 1998). However, there may also be other postsynaptic mechanisms in response to a chronic reduction of presynaptic transmitter release.

The neuromuscular junction of the nematode *Caenorhabditis elegans* comprises cholinergic and GABAergic MNs that directly innervate muscle. Moreover, cholinergic neurons that innervate muscles ventrally also synapse onto GABAergic neurons, innervating dorsal muscle, in order to enable body bending to occur; this topology also occurs in the inverse fashion, by different subsets of MNs (Stetina et al., 2006); see also **Fig. 9**). GABAergic feedback to cholinergic neurons is then mediated by GABA_B_ receptors (Dittman and Kaplan, 2008; Schultheis et al., 2011). Optogenetic stimulation experiments suggest that cholinergic MNs desensitize in response to repeated or prolonged stimulation, while GABAergic neurons appear to be potentiated (Liewald et al., 2008; Liu et al., 2009; Schultheis et al., 2011; Kittelmann et al., 2013), thus demonstrating some acute presynaptic plasticity of the MNs. Work from our lab highlighted an additional layer of regulation through neuropeptides (Steuer Costa et al., 2017). These neuropeptides were released in response to optogenetic cAMP generation and, in an autocrine fashion, caused a modulation of the ACh content of SVs. Another publication described the regulation of presynaptic cholinergic output when the function of acetylcholine esterase is inhibited, and this regulation involved neuropeptides that were released from a proprioceptive neuron, DVA (Hu et al., 2011). Furthermore, postsynaptic feedback mechanisms cause regulation of presynaptic transmission. One such mechanism is mediated through a microRNA (encoded by *mir-1*) that regulates postsynaptic nAChR expression but also presynaptic ACh release; the latter involves the extracellular cleavage of postsynaptic neurexin, that is released from the muscle surface in response to excess ACh signaling and inhibits presynaptic CaV2 channels and abundance of the SV protein RAB-3 (Simon et al., 2008; Hu et al., 2012; Tong et al., 2017). How the muscle may respond to altered ACh signaling (in this case, in response to increased ACh signaling / cholinergic agonists) to induce these changes is not well understood. However, it involves regulation of a transcription factor, MEF-2 (Simon et al., 2008). Whether muscle responds in a plastic way to reduced presynaptic ACh signaling is unknown.

As we have shown, cAMP signaling induced by optogenetics, using the photoactivated adenylyl cyclase bPAC in cholinergic neurons, causes an increase in the rate of SV fusion, but also enhances neuropeptide release. These neuropeptides act presynaptically to promote filling of acetylcholine into SVs via the vAChT (encoded by *unc-17* in *C. elegans*; (Steuer Costa et al., 2017). This was evident at the level of miniature postsynaptic currents (mPSCs), which exhibited increased mPSC rate and amplitude, and by the size of SVs as observed by electron microscopy, showing that bPAC stimulation rapidly induces an increase of SV diameter. In *unc-31* mutants, lacking the Ca^2+^ activator protein for secretion (CAPS) that is required for dense core vesicle (DCV) fusion and in which the release of neuropeptides is largely abolished, the increase of mPSC amplitudes was absent (Steuer Costa et al., 2017), and these animals had smaller SV diameters. Synaptic transmission at the NMJ is largely reduced in *unc-31* mutants, as both electrically or optogenetically evoked ePSCs are significantly smaller in *unc-31* mutants compared to wild type (Gracheva et al., 2007; Cornell et al., 2022). This is in line with the idea that SVs in these animals contain less ACh. The identity of the neuropeptides underlying this regulation at the NMJ is unclear, though NLP-12 neuropeptides, released from the proprioceptive neuron DVA (Li et al., 2006), were previously shown to affect cholinergic signaling at the NMJ, and to contribute to presynaptic plasticity (Hu et al., 2011). Animals which had been treated with the acetylcholine-esterase inhibitor aldicarb showed homeostatic upregulation of synaptic transmission that depended on the neuropeptide receptor CKR-2, expressed in cholinergic neurons. In recent years, single-cell RNAseq data identified the neuropeptides and neuropeptide receptors expressed in the MNs of *C. elegans*, as well as a functional analysis of cognate neuropeptide / neuropeptide receptor pairs (Taylor et al., 2021; Beets et al., 2023; Ripoll-Sanchez et al., 2023; Smith et al., 2024). Thus, the information of which peptides may be involved in NMJ plasticity is available, and we now wanted to functionally probe candidate neuropeptides. Furthermore, we wanted to analyze the consequence of the absence of neuropeptide signaling on the function of the NMJ.

Here, we identified several neuropeptides expressed in cholinergic and GABAergic MNs as being involved in cAMP-modulated cholinergic transmission, using behavioral assays and candidate mutants. We found NLP-9 and NLP-38 to be potentiators of cholinergic mPSC amplitude, and show an involvement of other neuropeptides and neuropeptide receptors. These neuropeptides depend on processing by two of the four proprotein convertases in *C. elegans*, AEX-5 and EGL-3, of which AEX-5 is specifically required in GABAergic neurons. The lack of these neuropeptides causes postsynaptic homeostatic upscaling of the ACh response in muscle. This was not affected through regulation of ACh receptors, but by upscaling the L-type VGCC EGL-19/CaV1. Lack of neuropeptide signaling induced the generation of larger action potentials (APs) in muscle, thus muscle cells upregulate their excitability to balance the reduced ACh release from presynaptic MNs.

## RESULTS

### UNC-31/CAPS is required in cholinergic and GABAergic neurons for normal synaptic transmission at the NMJ

To identify neuropeptides functionally involved in the cAMP regulation of transmission at the NMJ, we first tested different approaches that would qualify to demonstrate the involvement of a particular neuropeptide in NMJ regulation. To this end we used behavioral assays, combined with optogenetic bPAC stimulation of the cholinergic MNs, in wild type and *unc-31(n1304)* mutant animals. We assayed a) locomotion speed of crawling animals (Swierczek et al., 2011), b) body length as a readout for muscle tone (Liewald et al., 2008), and c) Ca^2+^ transients in postsynaptic muscle, using the genetically encoded fluorescent Ca^2+^ indicator RCaMP (Akerboom et al., 2013)(**Fig. 1A-F**). Acute activation of bPAC induces a significant speed increase in crawling animals (Steuer Costa et al., 2017) (**Fig. 1A, B**). The increased speed observed in wild type (wt) animals was robust, and significantly larger in *unc-31* mutants. It was further accompanied by mild body contraction (**Fig. 1C, D**), likely due to the coordinated activity of body wall muscles during locomotion induction. In this assay, *unc-31* mutants showed much reduced basal locomotion speed, as previously observed (Charlie et al., 2006), likely due to the general defect in neuropeptide signaling throughout the nervous system, though the phenotype may also be more specifically due to reduced ACh release at the NMJ of *unc-31* mutants. Thus, we expected a smaller increase of locomotion speed in response to bPAC activation in *unc-31* animals. Unexpectedly, however, they displayed a much more pronounced speed increase (almost reaching absolute speed levels of wt animals; **Fig. 1A, B**), as well as larger accompanying muscle contraction (**Fig. 1C, D**). This extent of the speed increase was surprising, given the fact that *unc-31* animals have much smaller evoked currents (Gracheva et al., 2007; Cornell et al., 2022) and smaller mPSCs. Possibly, compensatory changes increase the response of muscle to the reduced cholinergic transmission, a phenotype we have observed earlier for mutants affecting presynaptic transmitter release (Liewald et al., 2008). In line with the behavioral results, Ca^2+^ levels in muscles, following presynaptic bPAC photoactivation, elicited a more robust increase in *unc-31* mutants, as compared to wt animals (**Fig. 1E, F**). This indicates that reduced ACh signaling causes an increase in muscle excitability and/or Ca^2+^ influx. As an additional assay d), and to test whether postsynaptic muscles may have altered excitability, just as we had previously observed for mutants with reduced presynaptic ACh release (Liewald et al., 2008), we used direct optogenetic depolarization via channelrhodopsin-2 (ChR2) expressed in body wall muscle (BWM; *pmyo-3* promoter). Wild type animals showed body contraction by about 5 % of the body length in response to photostimulation (**Fig. 1G-J**). Indeed, *unc-31* mutants showed significantly enhanced body contraction of about 7.5 %.

**Figure 1.**
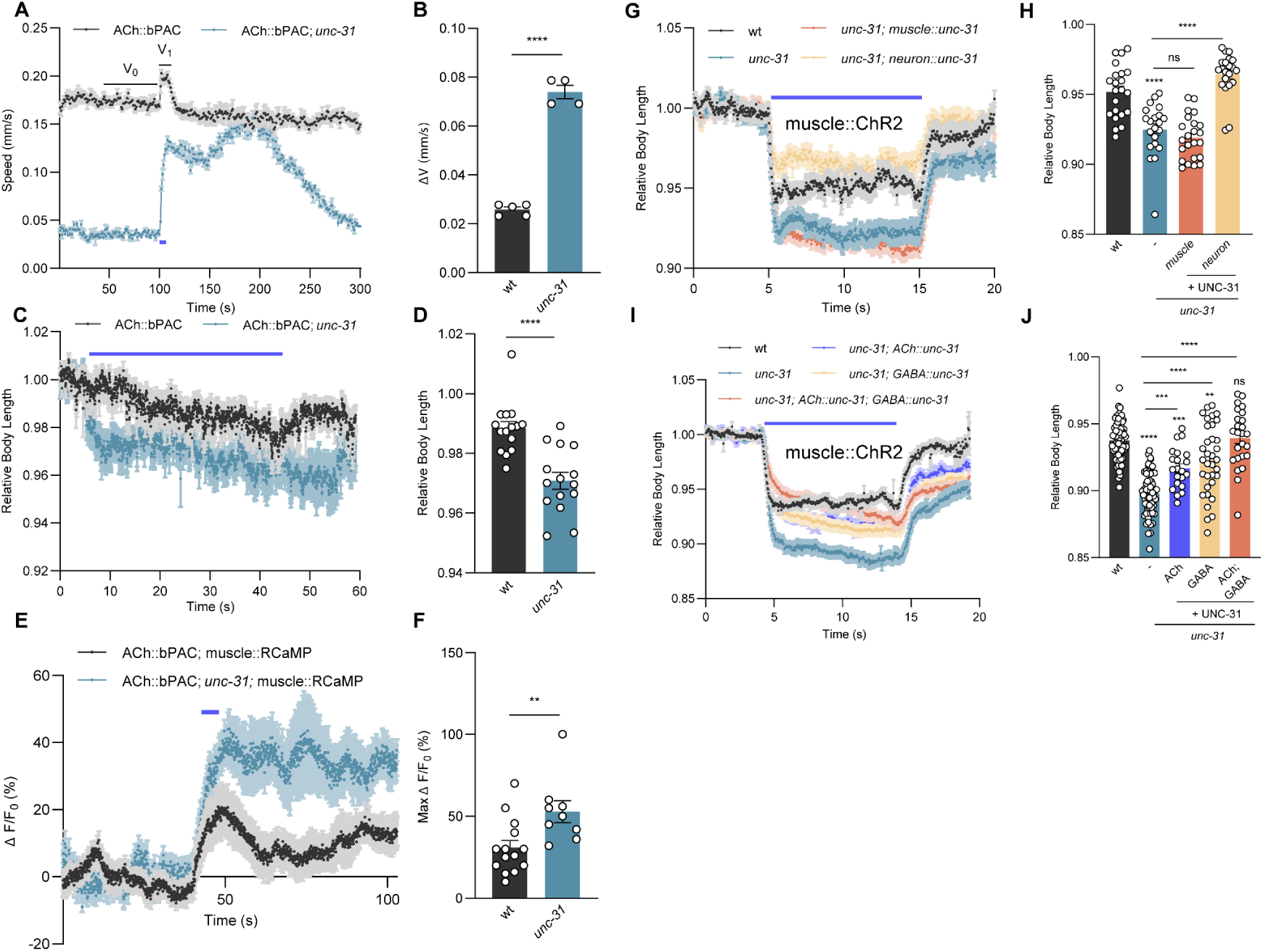
*unc-31* functions in cholinergic and GABAergic neurons to regulate NMJ synaptic transmission. **A, B)** Measurements of crawling speed of bPAC transgenic animals (using the *unc-17* promoter for cholinergic expression) on a multi-worm tracking device (MWT; Swierczek et al., 2011). Blue bar indicates 5 s period of blue light illumination. The average speed from 50-100 s was taken as basal speed V_0_ and from 100-120 s as V_1_. ΔV indicates the difference between V_0_ and V_1_. wt (n = 60-80, N = 5) and *unc-31(n1304)* mutants (n = 60-80, N = 4) were compared. An intensity of 70 µW/mm^2^ blue light was used for bPAC stimulation throughout this study. **C, D)** Measurements of (induced changes in) body length along the midline of single animals, before, during and after cholinergic bPAC activation. Absolute length values of worms (in pixels) were normalized to the averaged values before illumination (0-4 s, normalization was done for each animal) to obtain the presented relative body length. Blue bar indicates blue light illumination from 5 - 45 s. (wt, n = 15; *unc-31*, n = 15) **E, F)** Calcium imaging using the genetically encoded fluorescent indicator RCaMP in body wall muscle cells, increase evoked by cholinergic bPAC stimulation. RCaMP signal is normalized as ΔF/F_0_, where F_0_ means the average signal of first 40 s before blue light illumination, ΔF indicates the difference between the fluorescent signals F and F_0_. Max ΔF/F_0_ denotes the maximal signal observed after bPAC stimulation. Blue bar indicates blue light illumination from 45 - 50 s. (wt, n = 14; *unc-31*, n = 9) **G-J)** Measurements of body length induced by muscular ChR2 activation using 0.1 mW/mm^2^ blue light stimulation. *unc-31* rescue was performed by specifically expressing UNC-31 in muscles and neurons of *unc-31(n1304)* mutants (G-H; animal number tested in each group: n = 22, 21, 23, 21 from left to right columns, respectively), as well as in cholinergic or GABAergic neurons, and simultaneously in both neural types, respectively (I-J; animal number tested in each group: n = 62, 58, 21, 34, 25 from left to right columns). Blue bar indicates the 5 - 15 s blue light illumination. All data are presented as mean ± SEM. Statistical significance for two-group datasets and multiple-group datasets comparison was determined using unpaired t test and one-way ANOVA with Tukey-correction respectively. **, ***, and **** indicate p < 0.01, p < 0.001, and p < 0.0001, respectively.

As we have shown previously, neuropeptide signaling affects ACh release at the NMJ (Steuer Costa et al., 2017). Since UNC-31 affects the release of many if not most neuropeptides in *C. elegans*, and as different motor neuron classes at the NMJ feedback on each other (Stetina et al., 2006; Dittman and Kaplan, 2008; Schultheis et al., 2011; Zhen and Samuel, 2015), the source of the specific neuropeptides affecting cholinergic signaling may not be the cholinergic neurons. We thus sought to determine the site of action of UNC-31 in this context. The *C. elegans* motor circuit comprises cholinergic and GABAergic MNs (Stetina et al., 2006; Tolstenkov et al., 2018; Mulcahy et al., 2022). Cholinergic neurons form dyadic synapses with postsynaptic muscle and GABAergic neurons (White et al., 1986), and these innervate the opposite side of the body, thus evoking a body bend. Therefore, bPAC stimulation in cholinergic neurons also stimulates GABAergic neurons, and the observed behavioral effects, as well as the altered signaling in *unc-31* mutants, may have a GABAergic component. *unc-31* is primarily expressed in the ventral, dorsal, and sublateral nerve cords (Charlie et al., 2006; Gracheva et al., 2007; Cornell et al., 2022). We expressed UNC-31 in *unc-31* mutants specifically in body wall muscles (*pmyo-3* promoter) or pan-neuronally (*prab-3* promoter) (Charlie et al., 2006), and assessed body contraction in animals expressing ChR2 in muscle (**Fig. 1G, H**). As expected, UNC-31 function was rescued in neurons, but not in muscle. Since a focus of *unc-31* expression are cholinergic and GABAergic neurons (Charlie et al., 2006; Speese et al., 2007), we restored *unc-31* expression in both neuron types, respectively (**Fig. 1I, J**). UNC-31 expression in cholinergic and in GABAergic neurons could partially rescue the enhanced body contraction, but only when it was expressed in both cell types, a full rescue was observed. These results suggest that both cholinergic and GABAergic neuropeptides contribute to normal signaling at the NMJ, indicating complex neuromodulatory interactions.

### AEX-5 and EGL-3 proprotein convertases are required for NMJ neuropeptide signaling

Since UNC-31/CAPS is required for exocytosis of DCVs, which can contain different classes of neuropeptides as well as monoamines (Berwin et al., 1998; Speese et al., 2007; Hammarlund et al., 2008), we asked if different neuropeptide subclasses may be involved. The requirement of different neuropeptide precursor processing enzymes in different cell types may provide clues as to which neuropeptide(s) may be involved in NMJ regulation. Thus, we compared the muscle excitability of wt and mutant animals deficient in processing neuropeptide precursors. Those are processed by proprotein convertases (PCs) and carboxypeptidases to achieve mature and active neuropeptides (Canaff et al., 1999; Husson et al., 2006; Li and Kim, 2008). We tested the four known *C. elegans* PCs, three of which, *aex-5*, *bli-4*, and *kpc-1,* encode homologues of human PC1 with a cleavage sequence R-X-X-R, while the PC2 homologue KPC-1 is specific for cleavage following RR or KR sequences (Hung et al., 2014) (**Table S1**).

The *aex-5*/*kpc-3* mutants exhibited significantly enhanced muscle contraction compared to wt during ChR2 activation of BWMs, whereas *egl-3/kpc-2* mutants showed a slight, non-significant increase, and *kpc-1* and *bli-4/kpc-4* mutants showed unaltered contractions (**Fig. 2A, B**). *aex-5* mutants, however, were not as compromised as *unc-31* mutants in the same assay. With milder stimulation, *egl-3* mutants also exhibited enhanced muscle excitability (**Fig. S1A, B**), thus we investigated whether *aex-5* acts synergistically with *egl-3* in regulating NMJ signaling. *aex-5; egl-3* double mutants showed an additive effect, with body contraction that fully recapitulated the *unc-31* mutant phenotype (**Fig. 2C, D**). The enhanced muscle excitability of *aex-5* mutants could be rescued by restoring *aex-5* expression from its own promoter (**Fig. S1A, B**). This indicates that both enzymes jointly process a set of neuropeptides that are required for normal NMJ signaling, with *aex-5* playing a major role.

**Figure 2.**
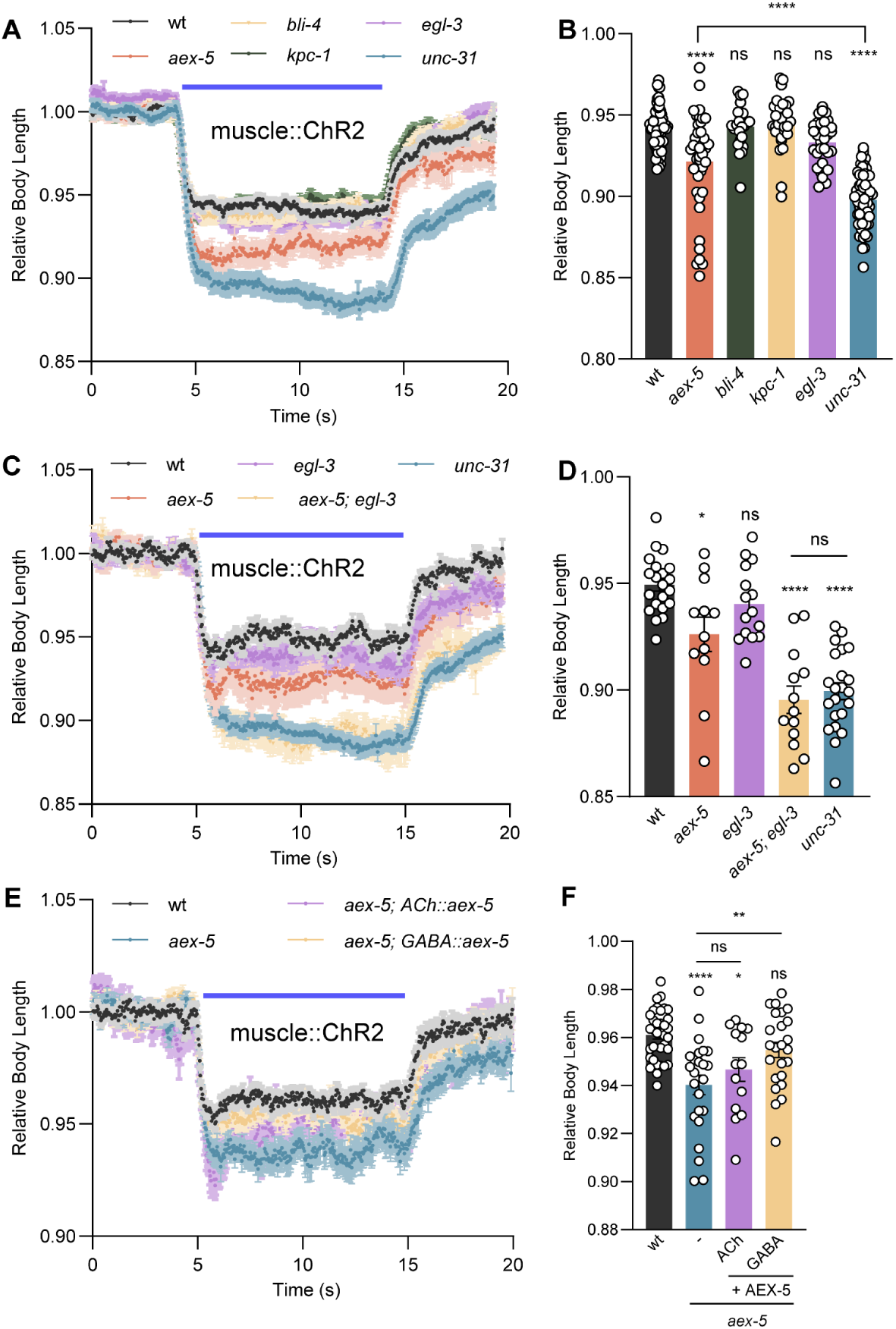
Proprotein convertases AEX-5 and EGL-3 are required for neuropeptidergic signaling at the NMJ. **A, B)** Measurements of body length induced by muscular ChR2 activation using 0.1 mW/mm^2^ blue light stimulation. Mean ± SEM and group data for time period during illumination. Single mutants *aex-5(sa23)* (n = 41), *bli-4(e937)* (n = 23)*, kpc-1(gk8)* (n = 31)*, egl-3(gk238)* (n = 32), and *unc-31(n1304)* (n = 58) were compared to wt (n = 60). **C, D)** as in (A, B) for mutants *aex-5* (n = 12), *egl-3* (n = 15), *unc-31* (n = 22), and *aex-5; egl-3* (n = 13) double knock out were compared to wt (n = 21). **E, F)** Measurements of body length 247 induced by muscular ChR2 activation using 65 µW/mm^2^ blue light stimulation. *aex-5* rescue was performed by specifically expressing AEX-5 in cholinergic and GABAergic neurons, animal number tested in each group: n = 35, 24, 14, 24 from left to right columns respectively. Blue bar indicates the 5 - 15 s blue light illumination. All data are presented as mean ± SEM. Statistical significance for multiple-group datasets comparison was determined using one-way ANOVA with Tukey-correction. *, **, and **** indicate p < 0.05, p < 0.01, and p < 0.0001, respectively.

*aex-5* was initially identified in a forward genetic screen for defecation phenotypes (Thomas, 1990; Husson et al., 2006). Later, it was reported that AEX-5 processes a peptide in the intestine that is released to regulate the defecation motor program (Mahoney et al., 2008). AEX-5 has also been reported to regulate AEX-1-dependent retrograde signaling at NMJs (Doi and Iwasaki, 2002), fat metabolism (Sheng et al., 2015), and pathogen avoidance (Singh and Aballay, 2019). We sought to determine whether AEX-5 also functions in cholinergic and GABAergic neurons. The muscle contraction phenotype of *aex-5* mutants was fully rescued by restoring AEX-5 expression in GABAergic, but not in cholinergic neurons (**Fig. 2E, F**). Furthermore, we detected the secretion of AEX-5 from GABAergic neurons, as it could be observed in coelomocytes, scavenger cells that capture released neuropeptides and PCs via bulk endocytosis (Sieburth et al., 2007; Mahoney et al., 2008) (**Fig. S1C**). These findings demonstrate that not only cholinergic, but also AEX-5 dependent GABAergic neuropeptide signaling is involved in regulating NMJ function.

### Functional screen for neuropeptides acting at the NMJ

We next wanted to identify neuropeptides affecting cholinergic signaling and/or the postsynaptic response to ACh release. The expression pattern of neuropeptides can be extracted from the available scRNAseq database CenGen (Taylor et al., 2021; Smith et al., 2024) (**Fig. S2**). Based on this data, we chose the most abundantly expressed neuropeptides in cholinergic MNs, i.e., FLP-6 (though expressed only in the VC neurons), FLP-9, FLP-18, NLP-9, NLP-13, NLP-15, NLP-21, and NLP-38. We also included neuropeptides expressed in GABAergic neurons that may likewise affect NMJ signaling and/or the excitation/inhibition balance at this synapse, i.e., FLP-15, but also FLP-9 and NLP-15, that are also expressed in cholinergic neurons. Last, we included NLP-12, expressed in the proprioceptive tail neuron DVA, as this was previously shown to affect cholinergic MNs and affects body bending; DVA itself is also cholinergic (Li et al., 2006; Hu et al., 2011). Interestingly, the GABAergic FLP-15 is also highly abundant in DVA. We used either the locomotion speed assay (bPAC in cholinergic neurons for this purpose, probing for neuron output; **Fig. 3A-C**), or the muscle contraction assay (ChR2 in BWMs, probing for altered excitability; **Fig. 3D, E**), to screen neuropeptides enriched in GABAergic neurons, since they may directly affect postsynaptic excitability other than via affecting cholinergic transmission.

**Figure 3.**
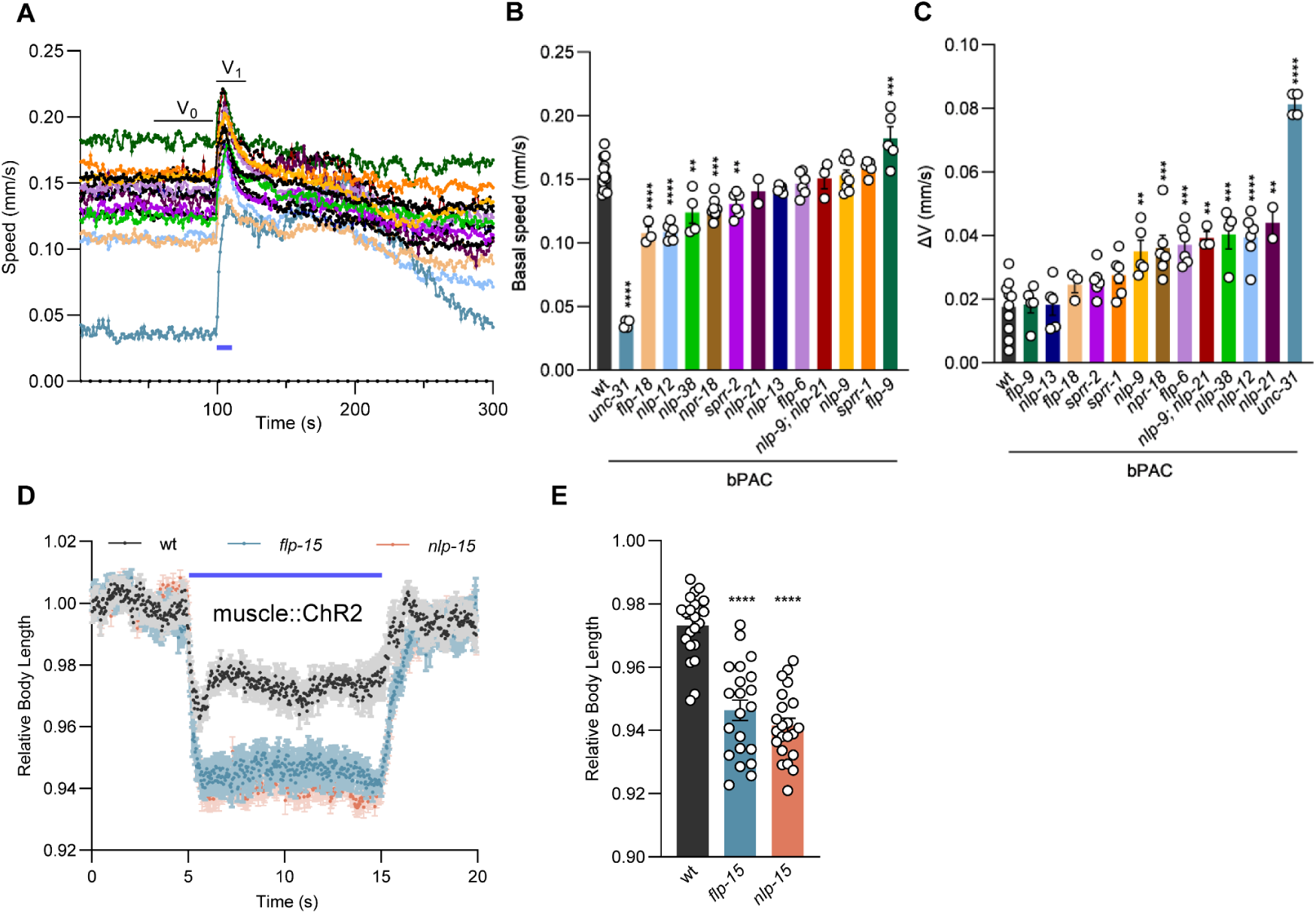
Candidate screen of neuropeptide / receptor genes affecting cholinergic transmission at the NMJ. **A-C)** Cholinergic-enriched neuropeptides were screened by measuring the crawling speed induced by cholinergic bPAC activation. Blue bar indicates the 5 s blue light illumination, at 70 µW/mm^2^. Mean speed traces (A), basal speed comparison (B), and speed increase comparison (ΔV = V_1_-V_0_) after bPAC activation (C) in animals of the indicated genotypes (wt, *flp-9, nlp-13, flp-18, sprr-2, sprr-1, nlp-9, npr-18, flp-6, nlp-9; nlp-21, nlp-38, nlp-12, nlp-21, and unc-31*, n = 60-80, N = 10, 5, 5, 3, 6, 6, 5, 6, 6, 3, 4, 6, 2, 4) are shown. **D, E)** Candidate GABAergic-enriched neuropeptides were assessed by measuring the body contraction induced by muscular ChR2 activation using 65 µW/mm^2^ blue light stimulation for 10 s. Two of the most abundant neuropeptides in GABAergic neurons *flp-15* (n = 21) and *nlp-15* (n = 22) were compared to wt (n = 22). All data are presented as mean ± SEM. Statistical significance for multiple-group datasets comparison was determined using one-way ANOVA with Tukey-correction. *, **, ***, and **** indicate p < 0.05, p < 0.01, p < 0.001, and p < 0.0001, respectively.

First, we analyzed the basal speed of mutants lacking neuropeptides in cholinergic neurons (**Fig. 3A, B**). *unc-31* mutants exhibit very slow locomotion. Likewise, also *nlp-12* mutants showed locomotion slower than wt. In addition, *nlp-38* and *flp-18* mutants were significantly slower, while *flp-9* mutants were faster than wt. Next, we assessed the speed increase following bPAC photoactivation (**Fig. 3A, C**). As expected, *nlp-12* mutants showed a significantly higher speed increase compared to wt, though not nearly as much as *unc-31* mutants (**Fig. 3A, B**). The speed increase in response to bPAC stimulation is very strong for *unc-31* mutants, and also *flp-6*, *nlp-9*, *nlp-21*, *nlp-38* and *nlp-9; nlp-21* double mutants showed a speed increase that was significantly more pronounced than in wt (**Fig. 3A, C**).

Of the neuropeptides highly expressed in GABAergic neurons (Taylor et al., 2021), *flp-15,* and *nlp-15* mutants exhibited enhanced muscle contraction in response to ChR2-mediated muscle depolarization (**Fig. 3D, E**), suggesting that the two neuropeptides contribute to regulation of NMJ transmission. In sum, the lack of several of the neuropeptides we tested partially phenocopies *unc-31* mutants. Possibly, combining all of these neuropeptide mutants might fully recapitulate the *unc-31* phenotype.

### The loss of NLP-9 in cholinergic neurons affects cholinergic signaling

The absence of a number of neuropeptides abundant in cholinergic neurons had significant effects, including FLP-6, NLP-9, NLP-12, NLP-21, and NLP-38. NLP-12 had been previously studied, NLP-21 is widely used as a marker for DCVs in cholinergic cells, and FLP-6 is expressed only in VC neurons. Therefore, and because a receptor for NLP-9 is known (NPR-18), we focused on testing the involvement of NLP-9 and NLP-38 neuropeptides in cholinergic transmission in more detail. First, we aimed to demonstrate the sufficiency of these neuropeptides for overall cholinergic output. The bPAC-induced speed increase that was exaggerated in *nlp-38* and *nlp-9* mutants was rescued by expression from their own promoters (**Fig. 4A, B**). Next, we tested muscle excitability more directly using ChR2 activation. *nlp-9* and *nlp-38* mutants showed enhanced muscle excitability compared to wt (**Fig. 4C, D**), and this was also observed in *nlp-9; nlp-21; nlp-38* triple mutants (**Fig. S3A, B**).

**Figure 4.**
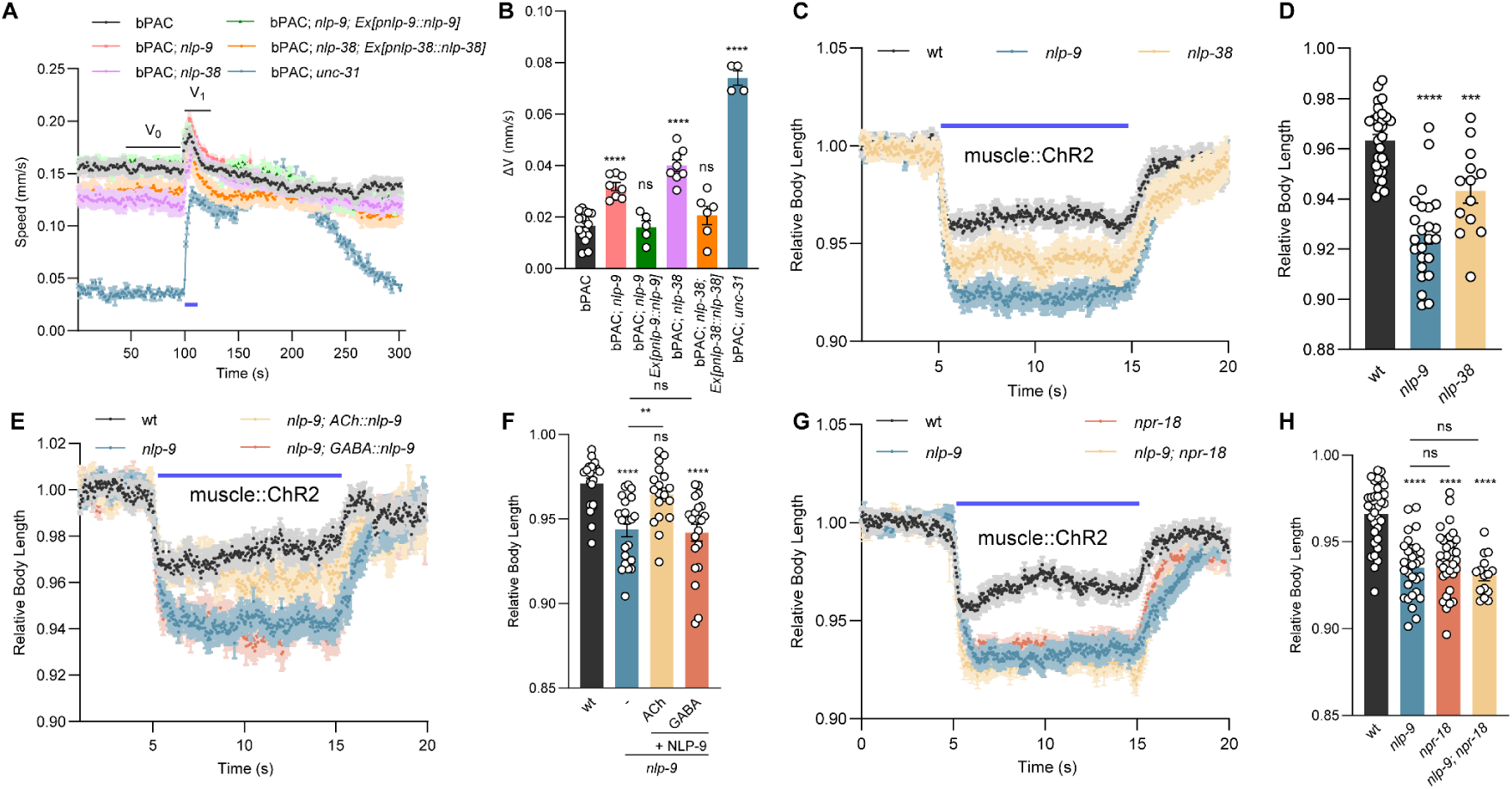
The loss of NLP-9 in cholinergic neurons induces enhanced postsynaptic excitability. **A, B)** Crawling speed induced by cholinergic bPAC activation was compared in the indicated genotypes. Blue bar indicates the 5 s blue light illumination (70 µW/mm^2^). *nlp-9* and *nlp-38* mutant rescues were achieved by specifically expressing NLP-9 and NLP-38 under their own promoters. Animal number tested in each group: n = 60-80, N = 16, 8, 5, 8, 6, 4 from left to right columns respectively. **C, D)** Body contraction induced by muscular ChR2 activation using 65 µW/mm^2^ blue light stimulation was compared in the indicated genotypes (wt, *nlp-9, nlp-38*, n = 29, 24, 13). **E, F)** Body contraction induced by muscular ChR2 activation using 65 µW/mm^2^ blue light stimulation was compared in the indicated genotypes. *nlp-9* mutants were rescued by specifically expressing NLP-9 in cholinergic and GABAergic neurons. Animal number tested in each group: n = 20, 22, 18, 22, from left to right columns respectively. **G, H)** Body contraction induced by muscular ChR2 activation using 65 µW/mm^2^ blue light stimulation was compared in the indicated genotypes (wt, *nlp-9, npr-18, nlp-9; npr-18*, n = 33, 30, 35, 15). Blue bar indicates the 5 - 15 s blue light illumination. All data are presented as mean ± SEM. Statistical significance for multiple-group datasets comparison was determined using one-way ANOVA with Tukey-correction. **, ***, and **** indicate p < 0.01, p < 0.001, and p < 0.0001, respectively.

A cell-type specific rescue experiment showed that NLP-9 is specifically required in cholinergic, but not GABAergic MNs (**Figs. 4E, F**; muscle excitability**; S3C, D**; bPAC induced speed increase); for NLP-38, we did not observe a cell-type specific rescue in an experiment testing bPAC induced speed increase (**Fig. S3E, F**). We assessed the expression pattern of both neuropeptide precursor genes. While NLP-9 showed a complete overlap with the expression pattern of the vesicular ACh transporter (vAChT) UNC-17 (**Fig. S4A**), NLP-38 showed a more restricted expression pattern to only a subset of cholinergic MNs (**Fig. S4B**). NPR-18 has been reported as the receptor for NLP-9 in modulating chemotaxis behaviors (Mills et al., 2012; Harris et al., 2014). Like *nlp-9* mutants, *npr-18* mutants exhibited enhanced muscle excitability (**Fig. 4G, H**), and *nlp-9; npr-18* double mutants did not show an additive effect, suggesting that they act in the same pathway. The reported expression pattern of NPR-18 is inconclusive, though: scRNAseq data suggests it is expressed in a small subset of cholinergic neurons, VC4 and 5; these cells, however, are innervating vulval muscles, not BWMs. In sum, *nlp-9* is required in cholinergic but not in GABAergic neurons for normal cholinergic evoked behavior, and is likely detected by NLP-18 receptors.

### NLP-9 neuropeptides are released from cholinergic neurons and affect ACh output, jointly with NLP-38

Next we wanted to assess the release of NLP-9 from cholinergic neurons. To this end, we expressed an mCherry-tagged version of the NLP-9 pro-protein in cholinergic neurons, and imaged the coelomocytes, i.e. cells that take up proteins and other materials from the body fluid (**Fig. 5A**). Fluorescent protein could be observed in these scavenger cells, and the levels of coelomocyte fluorescence were strongly reduced in an *unc-31* mutant background, confirming that the release of NLP-9 from cholinergic neurons is *unc-31* dependent (**Fig. 5B, C**). Furthermore, bPAC activation significantly increased the release of cholinergic NLP-9, supporting its role in modulation of cholinergic signaling (**Fig. 5B, C**).

**Figure 5.**
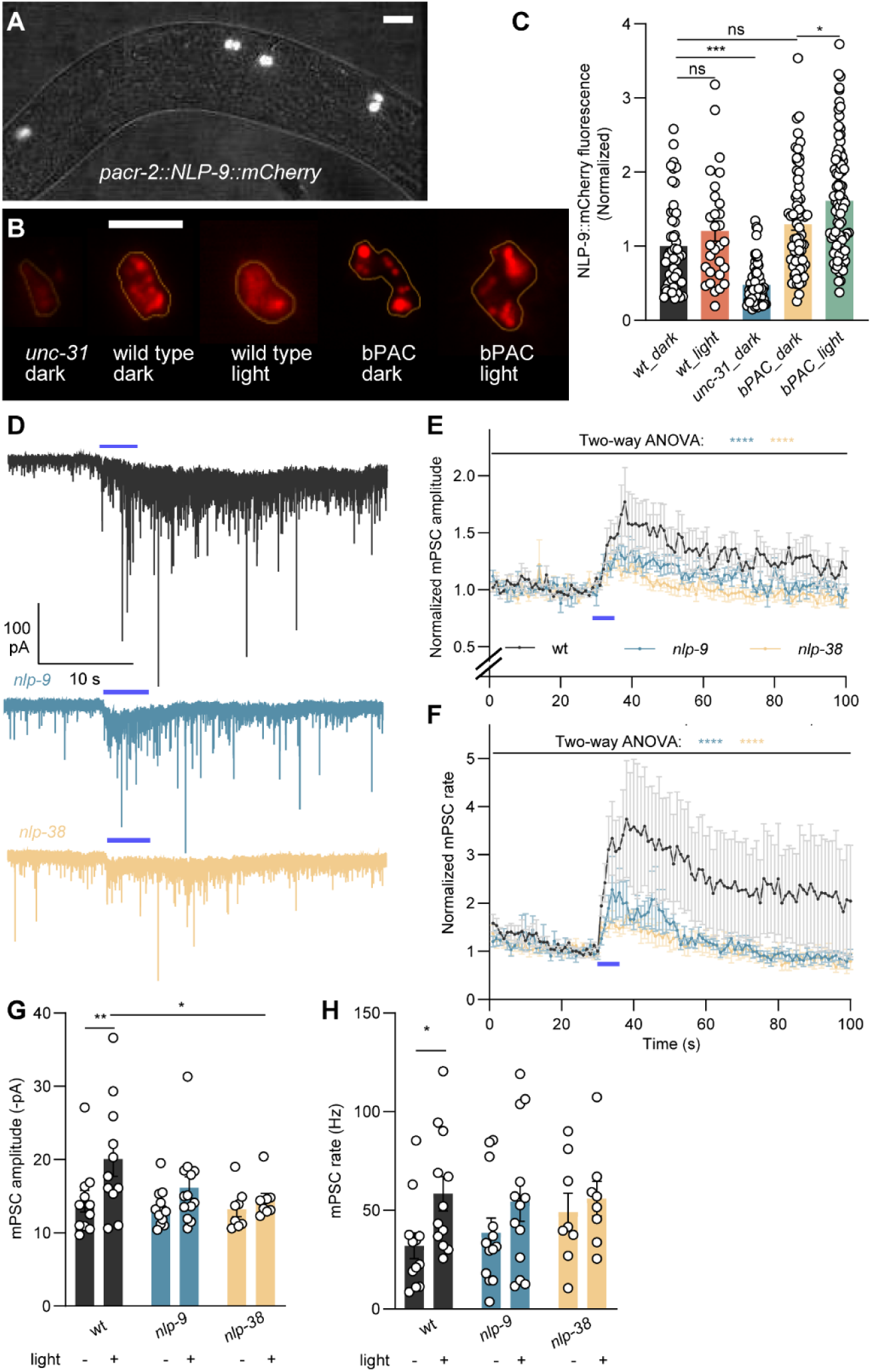
NLP-9 is released from choli-nergic neurons and affects ACh output, along with NLP-38. **A)** Fluorescence of endocytic scavenger cells (coelomocytes) resulting from secretion of mCherry-tagged NLP-9 proprotein, expressed in cholinergic MNs (using the *acr-2* promoter), into the body fluid. Scale bar 20 µm. **B)** Representative images showing NLP-9::mCherry fluorescence in coelomocytes from the indicated genotypes, and with or without blue light stimulation. Scale bar 10 µm. **C)** Fluorescence quantification of all coelomocytes in each animal, normalized to the signal in wt, is shown for the indicated genotypes and with or without blue light stimulation. Individual data points indicate single coelomocyte cells. Animal number tested n = 15, 8, 17, 19, 24, from left to right columns respectively. **D-H)** mPSCs recorded from body wall muscle of animals of the indicated genotypes. Representative traces of mPSC (D) from the indicated genotypes are shown. Blue bar indicates 5 s blue light stimulation. Normalized (to the 5 s before light stimulation) mean± SEM mPSC amplitudes (E) and mPSC rate (F) per second, averaged in 1 s bins. Summary data of mPSC amplitudes (G) and rate (H) are shown. Animal number n = 12, 13, 8 tested in wt, *nlp-9,* and *nlp-38,* respectively. All data are presented as mean ± SEM. Statistical significance comparison in (C) was determined using one-way ANOVA with Tukey-correction, and in (E-H) using two-way ANOVA with Tukey or Sidak test. *, **, ***, and **** indicate p < 0.05, p < 0.01, p < 0.001, and p < 0.0001, respectively.

Next, we analyzed *nlp-9* and *nlp-38* mutants lacking these neuropeptides by electrophysiology, i.e. whether they affect cholinergic transmission (**Fig. 5D**). Basal frequency and amplitude of mPSCs were not significantly altered in the two mutants (**Fig. 5G, H; S5A, B**), however, the amplitude increase of evoked PSCs was abolished in *nlp-9* and *nlp-38* mutants (**Fig. 5G, H**). Overall, the *nlp-9* and *nlp-38* mutants showed significantly different behavior (no amplitude increase and lower frequency increase than wt; **Fig. 5E, G**). Together, these findings establish NLP-9 and NLP-38 as regulators of cholinergic signaling at the NMJ, likely through controlling ACh content of SVs (Steuer Costa et al., 2017). NLP-9 (and likely, NLP-38) is released from cholinergic neurons in a cAMP-dependent manner. The absence of NLP-9 and NLP-38 (along with other neuropeptides) is thus likely to induce compensatory changes that, like in *unc-31* mutants, cause an enhanced behavioral response to (evoked) ACh release.

### Postsynaptic compensation in response to reduced cholinergic output

bPAC photo-stimulation and the induced cAMP signaling causes increased transmitter output, based on mobilization of SVs and their filling with additional ACh (Steuer Costa et al., 2017). If the loss of *unc-31* and neuropeptide signaling affects the extent of ACh output, one would expect that also the result of presynaptic depolarization would be affected in *unc-31* mutants. To investigate this, we stimulated cholinergic MNs using ChR2 photoactivation. Previous reports showed that ChR2-evoked (Cornell et al., 2022) and electrical stimulus-evoked EPSCs (Gracheva et al., 2007) in *unc-31* mutants are significantly decreased compared to wild type. However, counterintuitively, but consistent with the behavioral results induced by bPAC activation, we observed significantly enhanced muscle contraction in *unc-31* mutants after ChR2 activation, compared to wt (**Fig. 6A, B**). Also muscular Ca^2+^ increase was much larger in *unc-31* mutants (**Fig. 6C, D**). Based on these observations, we propose that reduced cholinergic transmission in *unc-31* mutants induces a postsynaptic compensatory mechanism in muscles, leading to enhanced muscle excitability and thus increased effects of evoked ACh release. As a consequence, the smaller presynaptic output in *unc-31* mutants could induce stronger muscle responses, to homeostatically maintain the synaptic strength.

**Figure 6.**
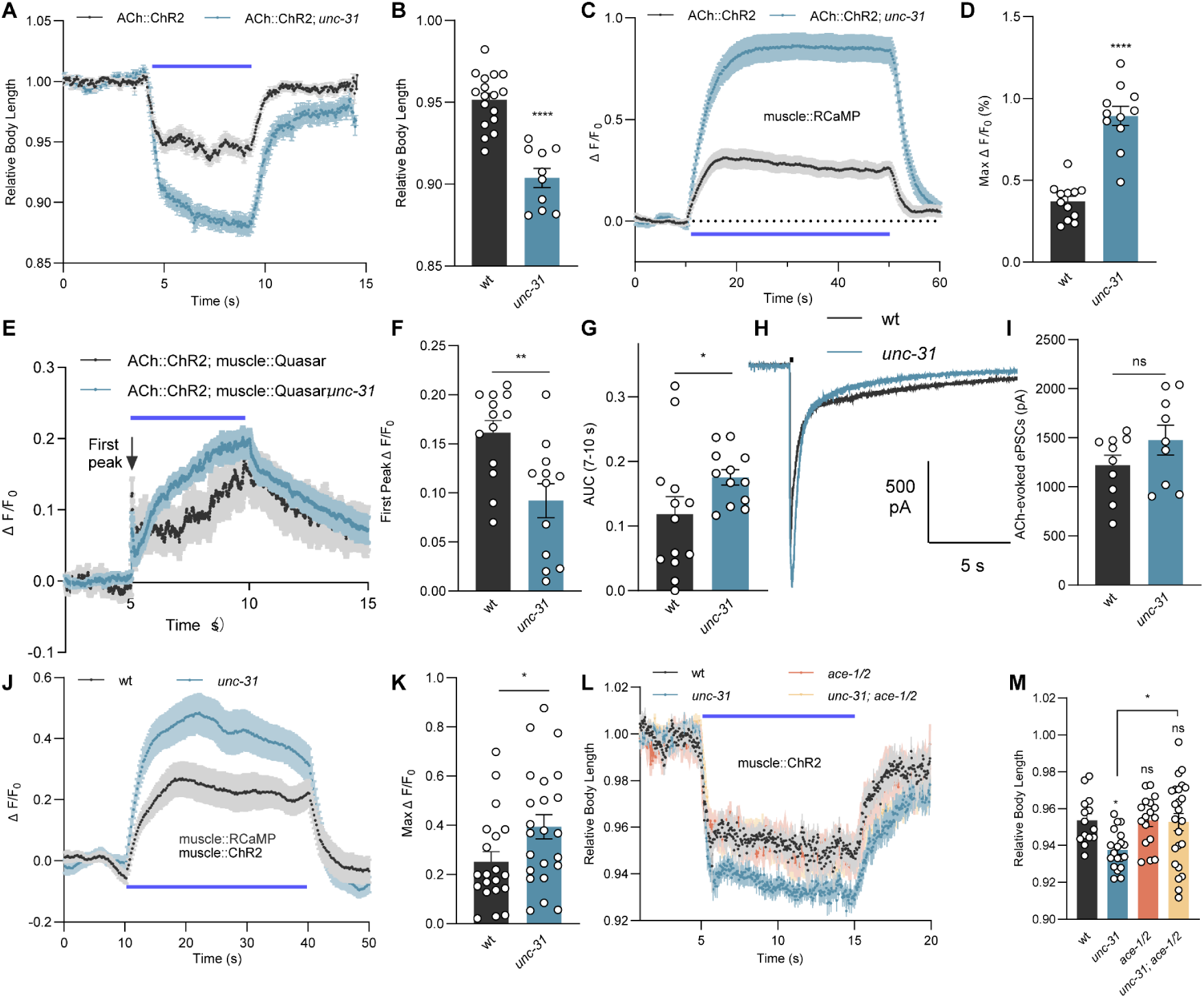
Reduced cholinergic transmission in *unc-31* mutants induces postsynaptic compensation. **A, B)** Body length change of wt (n = 17) and *unc-31* mutants (n = 10), induced by ChR2 activation in cholinergic neurons. **C, D)** RCaMP calcium imaging in BWM cells of wt (n = 12) and *unc-31* mutants (n = 11), signal increase evoked by cholinergic ChR2 stimulation. Max ΔF/F_0_ indicates the maximal signal observed after ChR2 stimulation. **E-G)** Voltage imaging using the fluorescent voltage indicator QuasAr, expressed in BWM cells of wt (n = 13) and *unc-31* mutants (n = 12), before, during and after depolarization evoked by ChR2 stimulation of cholinergic neurons. QuasAr fluorescent signal is normalized as ΔF/F_0_, where F_0_ is the average signal during the first 5 s before blue light illumination, and ΔF is the difference between the fluorescent signal F and F_0_. First peak ΔF/F_0_ is the initial maximal signal observed during ChR2 stimulation (F). AUC (7-10 s) indicates the area under the curve during the remaining stimulation period (G). **H, I)** ACh-evoked currents recorded from BWM cells of wt (n = 10) and *unc-31* mutants (n = 9). Representative traces (H) and group data of peak currents (I) are shown. **J, K)** RCaMP calcium imaging in BWM cells of wt (n = 20) and *unc-31* mutants (n = 23), signals evoked by muscular ChR2 stimulation. Max ΔF/F_0_, maximal signal after ChR2 stimulation. **L, M)** Body contraction induced by muscular ChR2 activation was compared in the indicated genotypes. n = 14, 19, 17, 24 animals tested of wt, *unc-31, ace-1; ace-2*, and *unc-31; ace-1; ace-2* animals, respectively. In all experiments except (H, I), ChR2 (in cholinergic neurons or muscles) was activated using 65 µW/mm^2^ blue light stimulation. All data are presented as mean ± SEM. Statistical significance for two-group datasets and multiple-group datasets comparison was determined using unpaired t test and one-way ANOVA with Tukey-correction, respectively. *, **, and **** indicate p < 0.05, p < 0.01, and p < 0.0001, respectively.

To test whether the functionally enhanced muscle excitability in *unc-31* mutants results from increased membrane excitability, we applied muscular voltage imaging, using the genetically encoded voltage indicator QuasAr (Azimi Hashemi et al., 2019; Bergs et al., 2023; **Fig. 6E**). In response to cholinergic ChR2 stimulation, voltage-dependent fluorescent signals in muscles showed a significantly reduced first peak in *unc-31* mutants, compared to wt (**Fig. 6E, F**). This was consistent with the reduced presynaptic ChR2- or electrical-evoked muscle EPSC currents (Gracheva et al., 2007; Cornell et al., 2022). However, during the remaining stimulation period (7-10 s), *unc-31* mutants showed increased voltage changes (**Fig. 6E, G**), indicating that the muscular plasma membrane becomes more depolarized during presynaptic stimulation, in line with the idea of enhanced muscle excitability.

The depolarization of body muscles involves the activation of two classes of nicotinic acetylcholine receptors (nAChRs): The heteromeric, levamisole-sensitive L-AChRs, and the homo-pentameric, nicotine-sensitive N-AChRs (Richmond and Jorgensen, 1999). Their activation by ACh triggers depolarization and Ca^2+^ entry via the voltage-gated calcium channel EGL-19 (Gao and Zhen, 2011; Laine et al., 2011; Liu et al., 2011). Possibly, also the T-type Ca^2+^ channel CCA-1 (Liu et al., 2011) contributes. We first asked whether the enhanced muscle excitability in *unc-31* mutants occurred at the level of nAChRs. Previously, we showed that the basal mPSC amplitudes were unaltered in *unc-31* mutants (Steuer Costa et al., 2017); **Fig. S6**), suggesting that the sensitivity of nAChRs in muscle cells to ACh remained unchanged. When we applied exogenous ACh to body wall muscles, the evoked postsynaptic currents (ePSCs) in wt and *unc-31* animals were indistinguishable (**Fig. 6H, I**), indicating that the expression and/or membrane delivery of nAChRs, as well as their functional properties, were unchanged. These results suggest that the compensatory increase in muscle excitability in *unc-31* animals is not induced at the level of nAChRs.

We thus reasoned that the enhanced muscle excitability could be caused downstream of nAChR activation. This would be in line with the experiments of direct muscle excitation using ChR2, thus probing changes in muscle excitability, independent of cholinergic signaling, and bypassing presynaptic influences. Muscular ChR2 activation evoked significantly increased muscle contraction (**Fig. 1G, H**) and Ca^2+^ increase (**Fig. 6J, K**) in *unc-31* mutants, compared to wt. Thus, muscle excitability appears to be enhanced in *unc-31* mutants, possibly in response to the reduced cholinergic transmission. To further validate the causality between the reduced cholinergic transmission and enhanced muscle excitability, we genetically manipulated presynaptic ACh levels in *unc-31* mutants. Acetylcholinesterases (AChEs) encoded by *ace-1* and *ace-2* break down ACh in the synaptic cleft. Consequently, AChE mutants should cause ACh accumulation in cholinergic synapses, and previous analyses demonstrated locomotion defects (Rand, 2007; Ringstad and Horvitz, 2008). Introducing *ace-1; ace-2* double mutations in the *unc-31* background indeed reverted the increased muscle excitability to wt level as the *unc-31*; *ace-1*; *ace-2* triple mutants showed a similar extent of muscle contraction as wt (**Fig. 6L, M**). However, *ace-1; ace-2* double mutants showed no significantly different muscle contraction compared to wild type, suggesting that an elevated presynaptic ACh level, contrary to its reduction, does not alter muscle excitability. This is in contrast to an earlier report describing upregulation of ACh output in response to pharmacological AChE blockade (Hu et al., 2011); however, these researchers used an acute assay, while in our experiment, mutants were used that may have undergone adaptation during development. Collectively, our results implicate that, in response to the decreased presynaptic ACh output in *unc-31* mutants, muscle excitability is homeostatically increased.

### Postsynaptic compensation involves EGL-19/Ca_V_1 L-type VGCCs

How is the muscle homeostasis responding to the loss of neuropeptidergic regulation achieved? As we had excluded the involvement of nAChRs in the homeostatic mechanisms (**Fig 6H, I**), we next assessed whether the machinery downstream of nAChRs was altered. The most immediate effects may be evoked by the L-type VGCC, encoded by *egl-19*. EGL-19 functions in the presynaptic terminal to regulate spontaneous release (Tong et al., 2017; Liu et al., 2018). In muscles, it regulates Ca^2+^-dependent action potentials and triggering of the ryanodine receptor UNC-68, as well as the muscle contractile apparatus (Gao and Zhen, 2011; Liu et al., 2011). Since *egl-19* null mutants are inviable, we acutely applied the specific antagonist nemadipine to inhibit the function of *egl-19* (Kwok et al., 2006). Nemadipine treatment significantly decreased the muscle contraction of both wt and *unc-31* mutants, elicited by muscle ChR2 activation (**Fig. 7A, B**).

**Figure 7.**
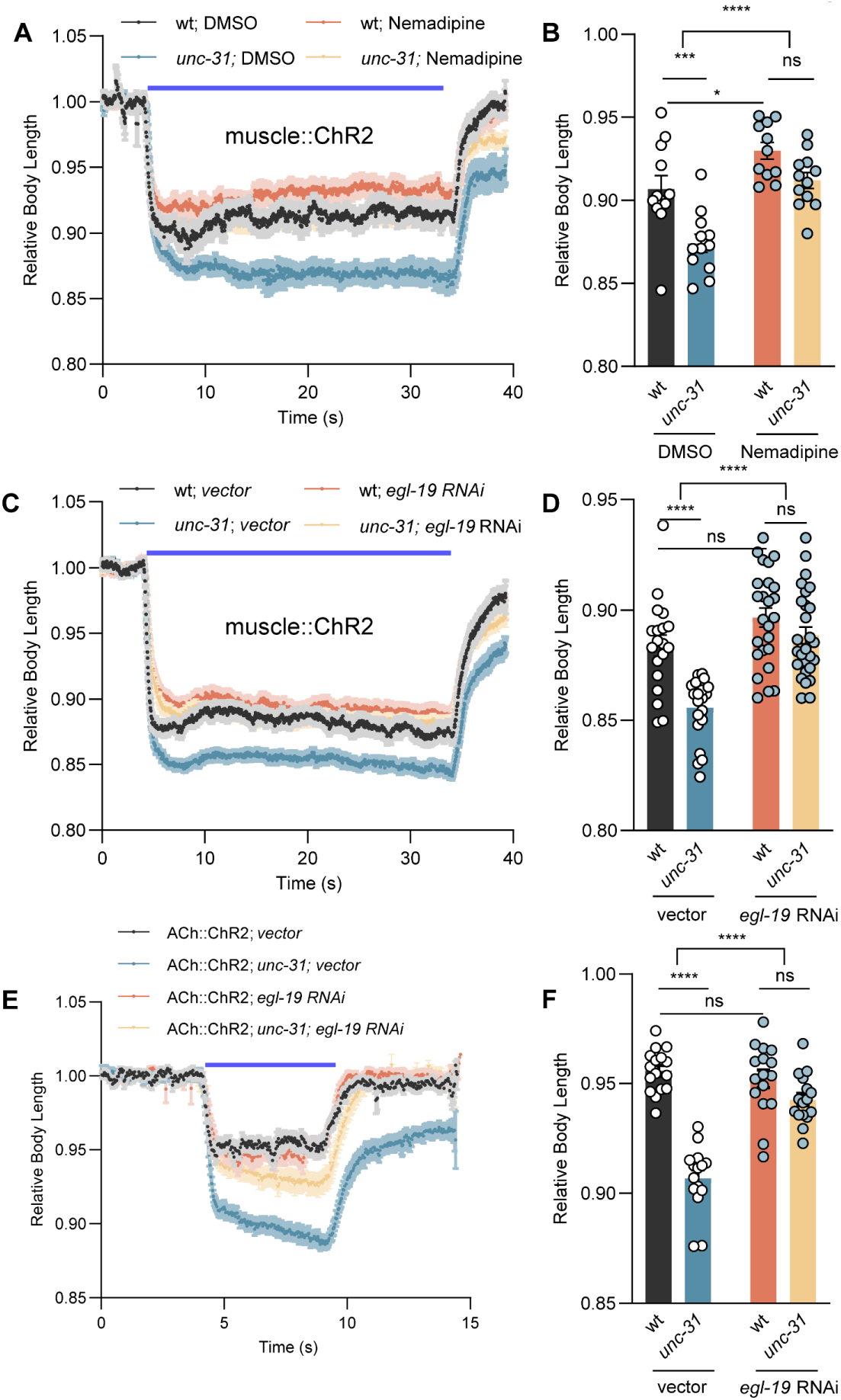
EGL-19 CaV1 mediates postsynaptic compensation of reduced presynaptic ACh release. **A-D)** Body contraction induced by muscular ChR2 activation using 1.1 mW/mm^2^ blue light stimulation was compared in the indicated genotypes and treatments. Relative body lengths after application of the EGL-19 CaV1 specific inhibitor nemadipine (A, B) and specific *egl-19* feeding RNAi (C, D) treatment are shown. DMSO-treated animals and animals fed the empty vector L4440 were used as controls. **E, F)** Body contraction induced by ChR2 activation in cholinergic neurons using 65 µW/mm^2^ blue light was compared in the indicated genotypes or RNAi treatments. Relative body lengths after specific *egl-19* RNAi treatment are shown. Number of animals tested in (B) n = 12, 12, 11, 12; (D) n = 19, 21, 25, 27; (F) n = 16, 15, 16, 16, from left to right, respectively. All data are presented as mean ± SEM. Statistical significance was determined using one-way ANOVA with Tukey-correction and two-way ANOVA with Sidak’s multiple comparisons test. *, ***, and **** indicate p < 0.05, p < 0.001, and p < 0.0001, respectively.

More importantly, after nemadipine treatment, *unc-31* mutants showed muscle contraction that was not significantly different from that of wt. We further confirmed the role of *egl-19* in regulating muscle excitability by (partial) knockdown of *egl-19 via* RNA interference (RNAi) (Fraser et al., 2000; Mariol and Segalat, 2001). Specific *egl-19* RNAi abolished the difference in muscle contraction between wt and *unc-31* mutants (**Fig. 7C, D**). Together, these results suggest that *egl-19* may be required for the enhanced muscle excitability in *unc-31* mutants, though this interpretation is complicated by the fact that EGL-19 is also a main mediator of muscle contraction. Additionally, nemadipine treatment also diminished the difference of muscle contraction between wt and *aex-5* mutants, implicating the involvement of *aex-5* in regulating muscle excitability (**Fig. S7**). Moreover, also when we stimulated the presynaptic cholinergic neuron by ChR2, the enhanced body contraction seen in *unc-31* mutants was decreased to wt level by *egl-19* RNAi (**Fig. 7E, F**). Collectively, these results suggest that the enhanced postsynaptic excitability induced by the loss of neuropeptidergic regulation may be mediated by muscular EGL-19.

### EGL-19 expression is upregulated in *unc-31* mutants and promotes postsynaptic excitability

To investigate how the EGL-19 channel may be regulated in response to the loss of neuropeptides, and as a means to achieve homeostatic compensation, we first looked at its expression levels in wt and *unc-31* mutants. Endogenous EGL-19 in muscle cells was labeled using the split mCherry complementary system (a gift from Joshua M. Kaplan; (Feng et al., 2019; Gao et al., 2022; Zhao et al., 2023). We observed a significant increase in the mCherry fluorescence in *unc-31* mutants, indicating that the expression level of EGL-19 is upregulated (**Fig. 8A, B**).

**Figure 8.**
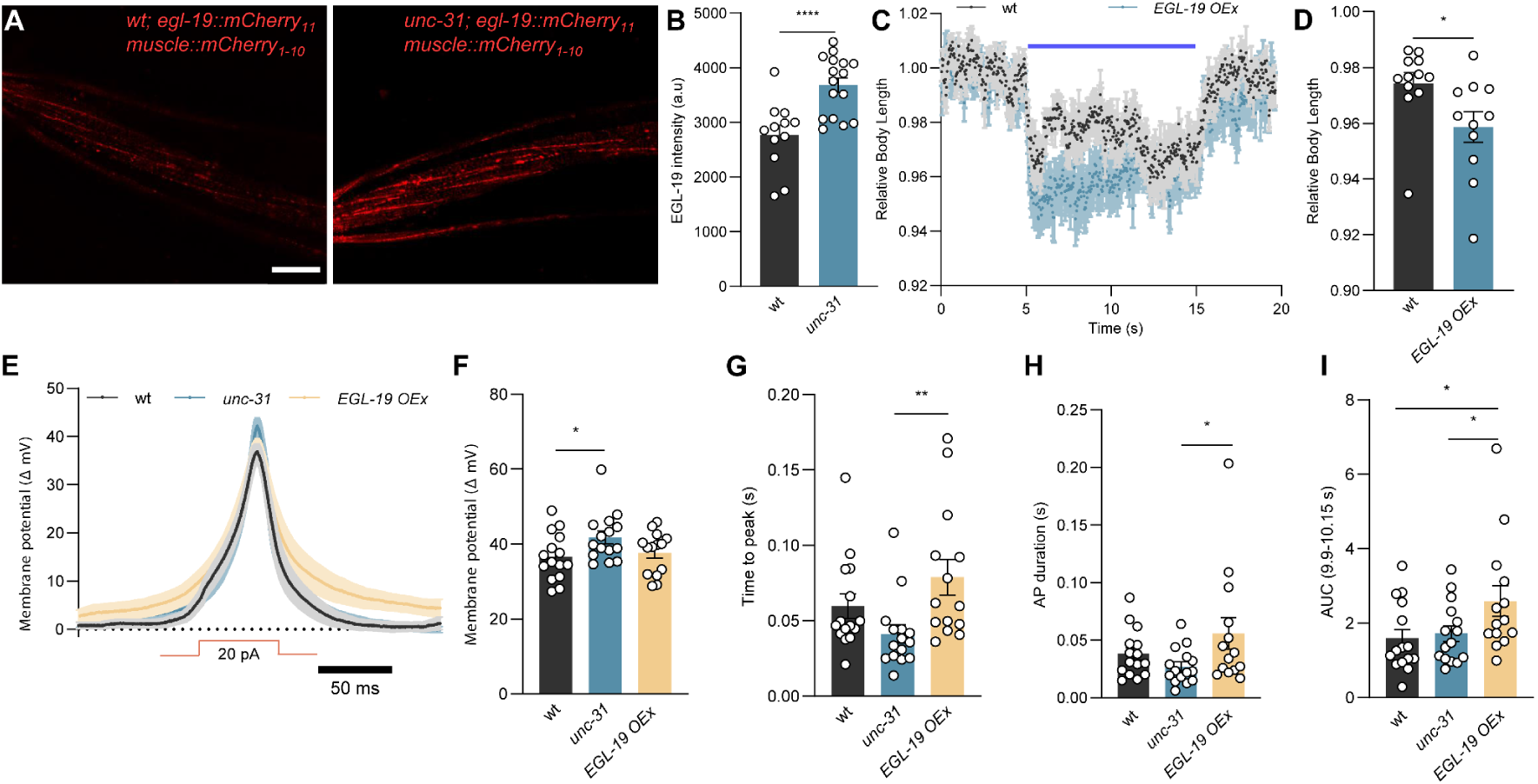
EGL-19 CaV1 is upregulated in *unc-31* mutants, leading to enhanced postsynaptic excitability. **A, B)** Endogenous expression levels of EGL-19 in muscle cells were compared in wt (n = 12) and *unc-31* mutants (n = 16). Representative images (A; Scale bar 20 µm) and summary data (B) are shown. **C, D)** Body contraction induced by muscular ChR2 activation using 65 µW/mm^2^ blue light stimulation. wt (n = 12) and muscle-specific EGL-19 overexpression (OEx) animals (n = 11) were compared. **E-I)** Current step-induced BWM membrane potential changes were compared in the indicated genotypes. Mean± SEM membrane potential changes (E) and summary data (F), mean time from current injection to reaching peak potentials (G), duration of the induced action potentials (H), and area under the curve (of the indicated time window 9.9-10.15 s) (I) are shown. Numbers of animal tested in (F-I) n = 15, 15, 14, from left to right columns respectively. All data are presented as mean ± SEM. Statistical significance for two-group datasets and multiple-group datasets comparison was determined using unpaired t-test, one-way ANOVA with uncorrected Fisher’s LSD test, or Kruskal-Wallis test with Dunn’s comparison, respectively. *, **, and **** indicate p < 0.05, p < 0.01, and p < 0.0001, respectively.

To further confirm EGL-19 regulates muscle excitability, we overexpressed EGL-19 in muscles and measured the body contraction induced by muscle ChR2 activation. Animals with EGL-19 overexpression displayed significantly stronger body contraction during ChR2 activation compared to wt, confirming that an elevated EGL-19 level induces higher muscle excitability (**Fig. 8C, D**). EGL-19 regulates action potentials (APs) in muscles (Gao and Zhen, 2011; Laine et al., 2011; Liu et al., 2011). If EGL-19 expression was upregulated in *unc-31* mutants, we would expect the APs to be potentiated as well. Consistent with this idea, in response to injecting current steps (20 pA, 50 ms) in patch-clamped muscle cells, the amplitude of induced APs was higher in *unc-31* mutants compared to wt (**Figs. 8E, F; S8**). However, in animals overexpressing EGL-19, this was not the case. Yet, EGL-19 overexpression caused the muscles to respond differently to the current step: The voltage increases started much delayed (**Figs. 8E, G; S8**) and the APs were longer lasting (**Figs. 8E, H; S8**). This led to a larger area under the curve for APs in EGL-19 overexpression animals (**Figs. 8E, I; S8**). Taken together, these results indicate that *egl-19* may play an important role in mediating postsynaptic compensation, which is triggered by the loss of presynaptic neuropeptidergic modulation. However, also other channels may contribute, like voltage-gated potassium channels that are required for the muscular AP downstroke (SHK-1, SLO-2; (Gao and Zhen, 2011; Liu et al., 2011), and the ER-resident ryanodine receptor, encoded by *unc-68*.

## DISCUSSION

Neuropeptidergic modulation of chemical synaptic transmission is likely to be a widely occurring phenomenon, and given the large variety of neuropeptide/neuropeptide-receptor systems, a lot of mechanisms may exist. Here, we have analyzed details of how the NMJ of the nematode *C. elegans* uses neuropeptides to regulate the output of ACh, and possibly GABA, to fine-tune transmission, but likely also to generally set the transmission output level. In the absence of neuropeptides, or when specific neuropeptide species, identified in this work, are missing, less transmitter output occurs which leads to an upregulation of postsynaptic excitability in the muscle cells. We showed that this occurs through the upregulation of the CaV1 VGCC EGL-19, and that the lack of neuropeptides causes the muscle to produce larger action potentials, likely to balance the missing cholinergic input. How the upregulation of CaV1 is achieved is currently unclear. It may involve previously discovered pathways for muscular regulation of excitability (Simon et al., 2008; Hu et al., 2012; Tong et al., 2017), though these involved the expression levels of nAChRs, which we did not find to be upregulated. Yet, gene expression mediated by the MEF-2 transcription factor might also affect EGL-19 expression. We propose that the neuropeptides work in an autocrine fashion to regulate presynaptic ACh output, however, it is possible that they also have postsynaptic effects. The latter appears unlikely, unless these peptides act inhibitory on gene expression, as it is the absence of neuropeptides that leads to CaV1 upregulation.

Both types of MNs are involved in neuropeptidergic regulation of the NMJ, as a complete rescue of *unc-31* deletion was achieved only if the CAPS protein was expressed in both cell types. Furthermore, the proprotein convertase AEX-5 rescue was only observed in GABAergic neurons. Thus, there is likely a complex interplay of neuropeptidergic regulation at the NMJ (Ripoll-Sanchez et al., 2023). This is evident from the number of neuropeptides as well as neuropeptide receptors that are expressed in the different motor neuron types (Taylor et al., 2021; Smith et al., 2024); deletion of several of those neuropeptides (NLP-9, -12, -15, -21, -38, FLP-6, -9, -15, -18), and of the NPR-18 receptor, affected (optogenetically induced) locomotion or muscle contraction. For NLP-9 and NLP-38, we could show by electrophysiology that there is an abolishment of the cAMP-induced increase of mPSC amplitudes. Several of these peptides may be part of autocrine feedback loops (NLP-9, -12, FLP-6, -9, -15, -18), as they are expressed along with their cognate receptors in MNs (Beets et al., 2023; Ripoll-Sanchez et al., 2023). However, often these peptides, or some of the peptides that are part of the individual precursor proteins, bind to a number of different receptors, expressed also in other parts of the nervous system, making interpretation of the findings less straightforward. We tested the effects of mutations in NPR-18 (putative receptor for NLP-9), SPRR-1, and SPRR-2 (receptor for NLP-38). Only NPR-18 had effects that were consistent with NLP-9. However, the expression pattern of NPR-18 (Taylor et al., 2021) shows expression in only the VC4 and VC5 neurons, that do not innervate muscle. SPRR-2 is also not obviously expressed in cholinergic neurons. The receptors for NLP-12, CKR-1 and CKR-2 (Hu et al., 2011; Beets et al., 2023), however, are expressed in almost all classes of cholinergic MNs. As we have shown previously, neuropeptides released from cholinergic neurons (this may include DVA, releasing NLP-12), affect the filling state of cholinergic SVs (Steuer Costa et al., 2017). Here, we identified these neuropeptides, thus one can now address how activation of neuropeptide receptors (and which ones) regulates filling of SVs via the vAChT UNC-17. Furthermore, we identified the requirement of proprotein convertases AEX-5 and EGL-3 for processing of the neuropeptides. Assessment of the expected cleavage sites in the neuropeptides we studied predicts a need for AEX-5 only in the case of NLP-12, while all others require EGL-3 (Table S1). Yet, NLP-12 is not predicted to be expressed by GABAergic neurons. Thus, more work is needed to clarify these details. However, cleavage specificity of individual neuropeptide precursors and need for a specific convertase should be confirmed by proteomics analyses of the respective mutants (Husson et al., 2006).

The postsynaptic compensation we observed in muscle in the absence of neuropeptides occurs via upregulation of the CaV1 VGCC, EGL-19. In line with this, *unc-31* mutants showed increased AP amplitudes. We explored if this could be mimicked by overexpressing EGL-19. While we did observe increased muscle contraction, just as in *unc-31* mutants, we did not observe an increased AP amplitude. However, APs were longer lasting, with a delayed onset and a larger AP integral, demonstrating that expression levels of EGL-19 can affect muscle excitability. Yet, neuropeptidergic regulation (or, the absence thereof) may induce additional modes of regulation of the CaV1 channel, not mimicked by its overexpression.

In sum, neuropeptides released from cholinergic and GABAergic neurons act in an autocrine fashion in cholinergic neurons, but also likely as a feedback loop from GABAergic to cholinergic neurons, to affect the output of ACh (**Fig. 9**). The lack of this signaling causes a reduction of SV filling with ACh (Steuer Costa et al., 2017) and thus reduced ACh output in *unc-31* / CAPS mutants. The reduced ACh output and thus reduced muscle depolarization causes a compensatory upregulation of EGL-19 CaV1 expression, leading to more Ca^2+^ entry despite less initial depolarizing signal from nAChRs. Whether such mechanisms are conserved in other animals will have to be demonstrated. In recent work we studied the larval zebrafish NMJ, in which optogenetically induced cAMP increase via bPAC also caused activation of locomotion behavior (Dill et al., 2024). Neuropeptide processing mutants lacking carboxypeptidase E, interestingly, also showed an increased response to the optogenetic stimulus. This, however, seemed to be affected on the level of nAChR expression in muscle cells, unlike in *C. elegans*, while the role of CaV1 channels had not been analyzed as yet.

**Figure 9.**
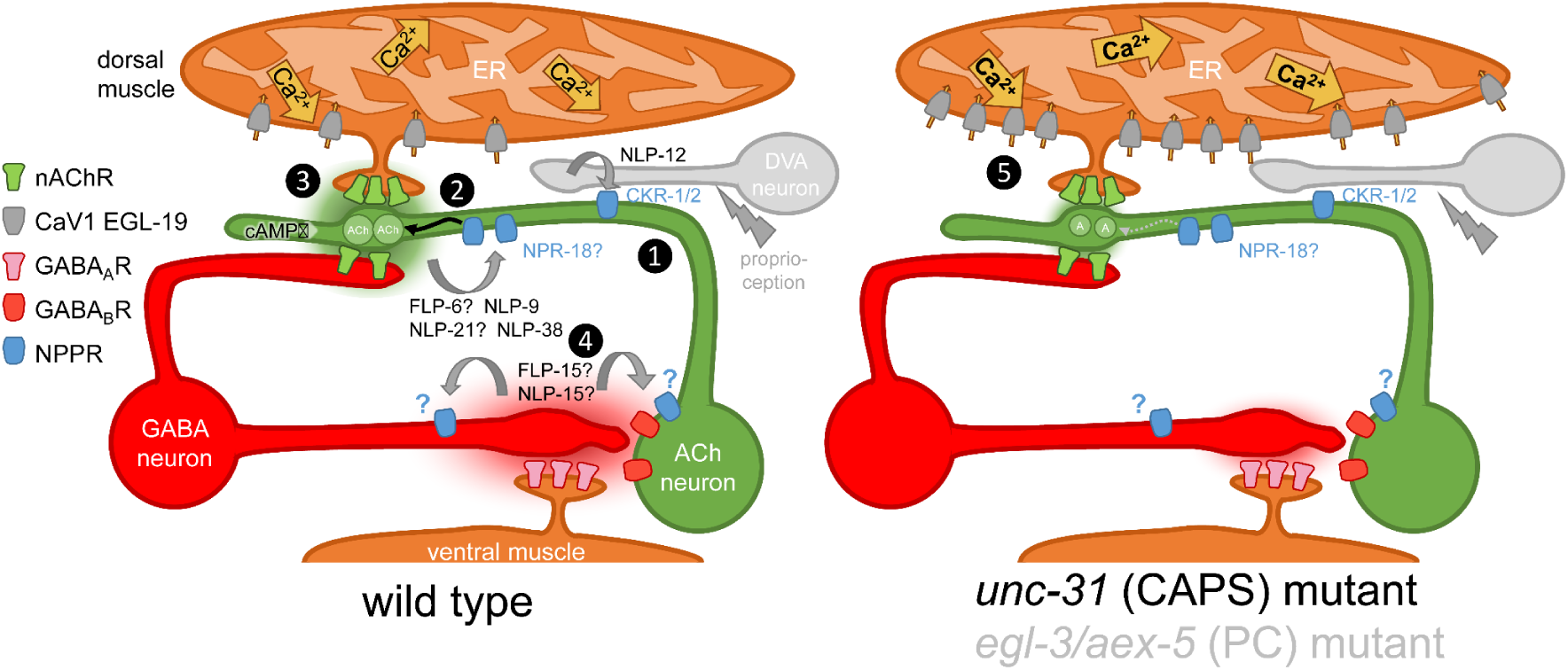
Model depicting findings made in this study. Neuromuscular junction of *C. elegans*, with cholinergic and GABAergic motor neuron, as labeled, innervating dorsal and ventral muscle, respectively. Dyadic synapse of cholinergic neuron to GABAergic neuron and muscle. Note that the same innervation pattern exists from different sets of MNs, but with inverted topology (not shown). Also shown is the proprioceptive neuron DVA, that innervates cholinergic neurons along the ventral nerve cord (drawn more dorsal, for simplicity). Relevant ion channels and receptors are indicated on the left (NPPR: neuropeptide receptor). In wild type (left), DVA releases NLP-12 neuropeptides onto cholinergic neurons, detected by CKR-1 and -2 receptors (1). (2) Cholinergic neuron releases ACh (green cloud) and neuropeptides FLP-6, NLP-9, NLP-21 and NLP-38, (likely) in an autocrine fashion, on NPR-18 and other, unknown neuropeptide receptors on cholinergic neuron. Activation of these receptors induces additional ACh filling of SVs (curved black arrow). (3) ACh release on muscle causes depolarization and Ca^2+^ influx into the cytosol, through CaV1 channels and from the ER (through UNC-68 ryanodine receptors, not shown); ACh release also activates GABAergic neuron. (4) GABA and neuropeptides FLP-15 and NLP-15 are released on muscle and cholinergic neurons, and are detected by GABA_A_ (muscle) and GABA_B_ receptors (cholinergic neuron), as well as by unknown neuropeptide receptors. In *unc-31* (CAPS) or *egl-3; aex-5* (proprotein convertases, PCs) double mutants (right half), no mature neuropeptides are released or formed, thus the signaling steps induced by neuropeptides are absent. As a consequence, (5) less ACh is filled into SVs and released from cholinergic neuron. In response, compensatory upregulation ensures homeostatic NMJ signaling by providing more Ca^2+^ influx and thus depolarization of muscle cell.

## Materials and Methods

### Worms and maintenance

*C. elegans* strains were cultivated at 20°C, on nematode growth media (NGM) plates seeded with *Escherichia coli* strain OP50 as previously described (Brenner, 1974).

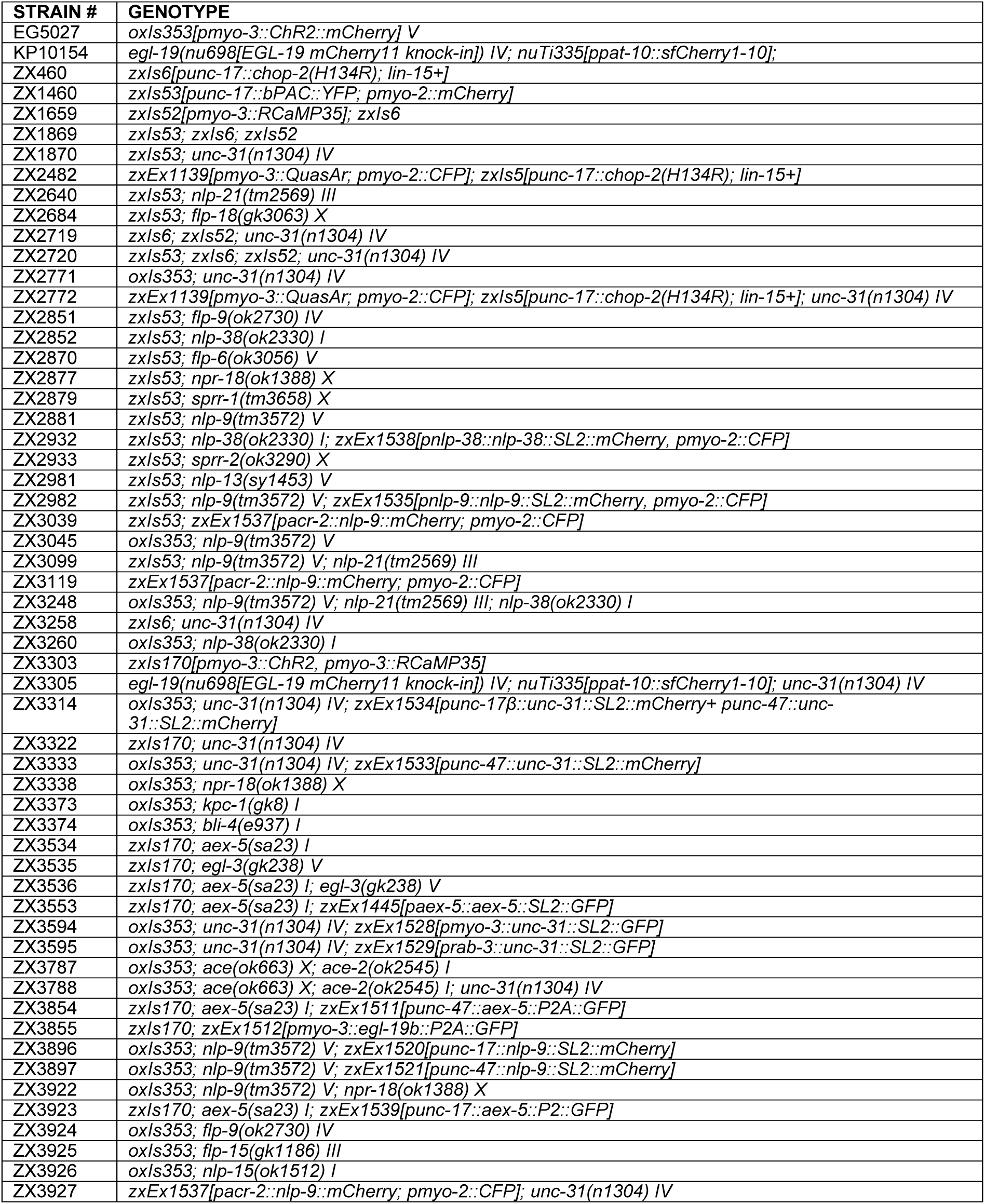

### Molecular biology

Plasmids generated or used in this work are listed in **Table S2**, and further described below:

### Rescue Constructs

*unc-31* cDNA was amplified by PCR from KG#121, a gift from Kenneth Miller (Addgene plasmid # 110879; http://n2t.net/addgene:110879; RRID: Addgene_110879), and inserted into expression vectors containing the *rab-3* promoter (pan-neuronal rescue), the *unc-17* promoter (cholinergic rescue), the *unc-47* promoter (GABAergic rescue), and the *myo-3* promoter (muscle rescue) respectively.

*aex-5, nlp-9* and *nlp-38* genomic sequence were amplified by PCR from N2 genomic DNA and inserted into expression vectors respectively.

Genomic fragments as promoters, upstream of the start codon of *aex-5* (2 kb), *nlp-9* (2.8 kb), and *nlp-38* (2 kb), were amplified by PCR from N2 genomic DNA.

### egl-19 RNAi

A 1.5 kb genomic fragment of *egl-19* (Mariol and Segalat, 2001) was amplified from N2 genome and inserted into plasmid L4440 by restriction digest with *HindIII* and *NheI*. The construct was transformed in *E. coli* strain HT115 (Timmons and Fire, 1998). dsRNA induction was performed as previously described (Wabnig et al., 2015).

### *egl-19* overexpression

*egl-19b* cDNA was amplified by PCR from KP#2460 (kindly provided by Joshua M. Kaplan). The cDNA was inserted into expression vector that contains the muscular expression promoter *myo-3*.

### Transgenes and Germline Transformation

Transgenic strains were generated by microinjection of various DNA constructs with a co-injection marker (pmyo-2::mCherry (2 ng/µl), pmyo-2::CFP (2 ng/µl), or pttx-3::mScarlet (30 ng/µl). Integrated strains were obtained by histone-miniSOG induced mutagenesis (Noma and Jin, 2015), followed by outcrossing against N2 at least 4 times.

### Behavioral assays

Crawling speed assay was performed in a multiworm tracker (MWT) platform (Swierczek et al., 2011), illuminated by LED modules ALUSTAR (Ledxon GmbH, Geisenhausen, Germany) at 70 µW/mm^2^, and as previously described (Vettkotter et al., 2022; Aoki et al., 2023). Due to the dark activity of bPAC, all experiments dealing with bPAC-expression strains were performed in a dark room, and animals were manipulated under red-filtered light (Steuer Costa et al., 2017). An infrared back light on the MWT was made with WEPIR3-S1 IR Power LED Star infrared (850 nm) 3W (Winger Electronics GmbH & Co. KG, Dessau, Germany) powered by LCM-40 constant current LED driver and regulated with a potentiometer (Vishay Intertechnology, Inc., Malvern, PA, USA) to avoid bPAC pre-activation. To synchronize worms, around 15 gravid animals were transferred to OP50-seeded NGM plates and allowed to lay eggs for 6-8 hours, and then removed. After 3-4 days, around 80 day-1 adults were collected using M9 buffer and transferred onto plain NGM plates for measurements.

Body contraction assay was performed in a single worm tracker, illuminated with blue light from a 50 W HBO mercury lamp filtered through a GFP excitation filter (450-490 nm) at 1.1, 0.1, and 65 µW/mm^2^ intensities as indicated in respective figure legends, under a 10x objective in a Zeiss Axiovert 40 microscope (Zeiss, Germany). L4 larvae were transferred to freshly seeded ATR plates (seeded with 300 µl OP50-1 culture, mixed with 0.6 µl of 100 mM ATR stock dissolved in ethanol) and single day-1 adults were placed onto plain NGM plates for measurements. The body length was determined as previously described (Seidenthal et al., 2022).

Animals were kept in darkness for 15 min before all recordings. The measurements were repeated on 2-3 different experimental days.

### Fluorescence imaging

For calcium imaging, combined with optogenetic stimulation, RCaMP35-(Akerboom et al., 2013) and ChR2-expressing animals were mounted on glass slides and immobilized with polystyrene beads (0.1 µm diameter, at 2.5% w/v, Sigma-Aldrich). Images were taken with a Zeiss Observer Z1 compound microscope equipped with a Kinetix22 (Teledyne Photometrics, Tucson, AZ, USA) camera, Prior Lumen LEDs (Prior Scientific, Cambridge, UK), and a RCaMP filter cube (a dual band excitation filter 479/585 nm, a 647 nm emission filter, and a 605 nm bean splitter, F74-423, F37-647, and F38-605, respectively; AHF Analysentechnik, Germany), with a 20x objective with illumination light at 470 nm to activate ChR2 at defined period, with excitation light at 590 nm for RCaMP signal recording.

Voltage imaging was performed as previously described (Azimi Hashemi et al., 2019). Briefly, QuasAr-expressing animals were immobilized with polystyrene beads and imaged on a Zeiss Axio Observer Z1 microscope equipped with 40× oil immersion objective (Zeiss EC Plan-NEOFLUAR ×40/N.A. 1.3, Oil DIC ∞/0.17), a laser beam splitter (HC BS R594 lambda/2 PV flat, AHF Analysentechnik), and a galilean beam expander (BE02-05-A, Thorlabs). Voltage-dependent fluorescence of QuasAr was excited with a 637 nm laser (OBIS FP 637LX, Coherent) at 1.8 W/mm^2^ and imaged at 700 nm, while ChR2 was activated by a monochromator (Polychrome V) at 300 μW/mm^2^. The Coelomocyte assay was performed as previously described (Sieburth et al., 2007). Briefly, NLP-9::mCherry fusion protein-expressing animals were mounted on glass slides and immobilized with tetramisole. Images were taken with Zeiss Observer Z1 with 40x objective with 590 nm illumination with RFP filter cube (ex. 580/23 nm, em. 625/15 nm). All coelomocytes within each animal were imaged for quantification. For bPAC strains, worms were illuminated with blue light at 35 µW/mm^2^ for 15 min prior to imaging.

Images of representative expression patterns of *nlp-9 and nlp-38* reporter constructs, as well as *EGL-19::mCherry* images for quantification, were acquired with a Zeiss LSM780 microscope with 63x oil objective (Plan-Apochromat 63x/1.4 Oil DIC) at 488 nm and 543 nm for GFP and mCherry excitation respectively.

Image were subjected to background subtraction and quantification conducted with Fiji (Schindelin et al., 2012).

### Electrophysiology

Electrophysiological recordings of body muscle cells were done in dissected adult worms as previously described (Liewald et al., 2008). Animals were immobilized with Histoacryl L glue (B. Braun Surgical, Spain) and a lateral incision was made to access neuromuscular junctions (NMJs) along the anterior ventral nerve cord. The basement membrane overlying body wall muscles was enzymatically removed by 0.5 mg/ml collagenase for 10 s (C5138, Sigma-Aldrich, Germany). Integrity of BWMs and nerve cord was visually examined via DIC microscopy. Recordings from BWMs were acquired in whole-cell patch-clamp mode at 20-22 °C using an EPC-10 amplifier equipped with Patchmaster software (HEKA, Germany). The head stage was connected to a standard HEKA pipette holder for fire-polished borosilicate pipettes (1B100F-4, Worcester Polytechnic Institute, USA) of 4–10 MΩ resistance. The extracellular bath solution consisted of 150 mM NaCl, 5 mM KCl, 5 mM CaCl_2_,1 mM MgCl_2_,10 mM glucose, 5 mM sucrose, and 15 mM HEPES, pH 7.3, with NaOH, ∼330 mOsm. The internal/patch pipette solution consisted of K-gluconate 115 mM, KCl 25 mM, CaCl_2_ 0.1 mM, MgCl_2_ 5 mM, BAPTA 1 mM, HEPES 10 mM, Na_2_ATP 5 mM, Na_2_GTP 0.5 mM, cAMP 0.5 mM, and cGMP 0.5 mM, pH 7.2, with KOH, ∼320 mOsm.

Voltage clamp experiments were conducted at a holding potential of -60 mV. Light activation was performed using an LED lamp (KSL-70, Rapp OptoElectronic, Hamburg, Germany; 470 nm, 8 mW/mm²) and controlled by the Patchmaster software. Puff-application (80 ms) of ACh (500 μM in CRG) to record ACh-evoked ePSCs were performed using a Picospritzer III (Parker, USA). Subsequent analysis and graphing was performed using Patchmaster, and Origin (Originlabs). Analysis of mPSCs was conducted with MiniAnalysis (Synaptosoft, Decatur,GA, USA, version 6.0.7). in body wall muscle cells were recorded in current clamp mode. Membrane potential and action potentials in BWM cells were recorded in current clamp mode. An additional current pulse of +20 pA (50 ms) was injected at 10.01 s via the Patchmaster software for 50 ms. For data analysis, voltage traces were aligned to the peak of the AP, and replotted, thus appear shifted relative to the current pulse.

### Statistical analysis

All quantitative data are shown as mean ± s.e.m. n indicates number of animals, and N indicates biological replicates / measurement sessions with distinct populations of animals, unless otherwise noted. Significance between data sets after two-tailed Student’s t-test (two groups comparison), one-way AVOVA (multiple groups comparison), with Bonferroni’s or Tukey’s multiple comparison test, or Fisher test, or two-way ANOVA (multiple groups comparison), were performed as indicated in each figure legend. Data were analyzed and plotted in GraphPad Prism (GraphPad Software, version 8.02) or OriginPro 2024 (64-bit; SR1 10.1.0.178; Origin Lab Corporation).

## Data and reagent availability

All reagents and data are available from the corresponding author.

## Author contributions

**Conceptualization:** J.S., W.S.C., A.G.

**Methodology:** J.S., J.F.L.

**Validation:** J.S., J.F.L., A.G.

**Formal analysis:** J.S., J.F.L., A.G.

**Investigation:** J.S., J.F.L., C.S., J.G., M.S.D., W.G.

**Data Curation:** J.S., J.F.L., A.G.

**Writing - Original Draft:** J.S., A.G.

**Writing - Review & Editing:** J.S., J.F.L., A.G.

**Visualization:** J.S., J.F.L., A.G.

**Supervision:** J.S., A.G.

**Project administration:** A.G.

**Funding acquisition:** A.G.

## Acknowledgements

We would like to express our gratitude to members of the Gottschalk lab in supporting this research. We are particularly indebted to Franziska Baumbach and Katharina Kuhlmeier, for excellent technical support. We thank Kenneth Miller, Stephen Nurrish and Joshua Kaplan for reagents and strains. Some strains were obtained from the *Caenorhabditis* Genetics Center (CGC), which is funded by the NIH Office of Research Infrastructure Programs (P40 OD010440). Other strains were obtained from the Japanese National Bioresource Project for the nematode, funded by the Japan Agency for Medical Research and Development (AMED). This work was funded by the Deutsche Forschungsgemeinschaft (DFG), CRC1080 project B02, and by core support from Goethe University, both to A.G.

## Supplementary Figures Legends

**Figure S1.**
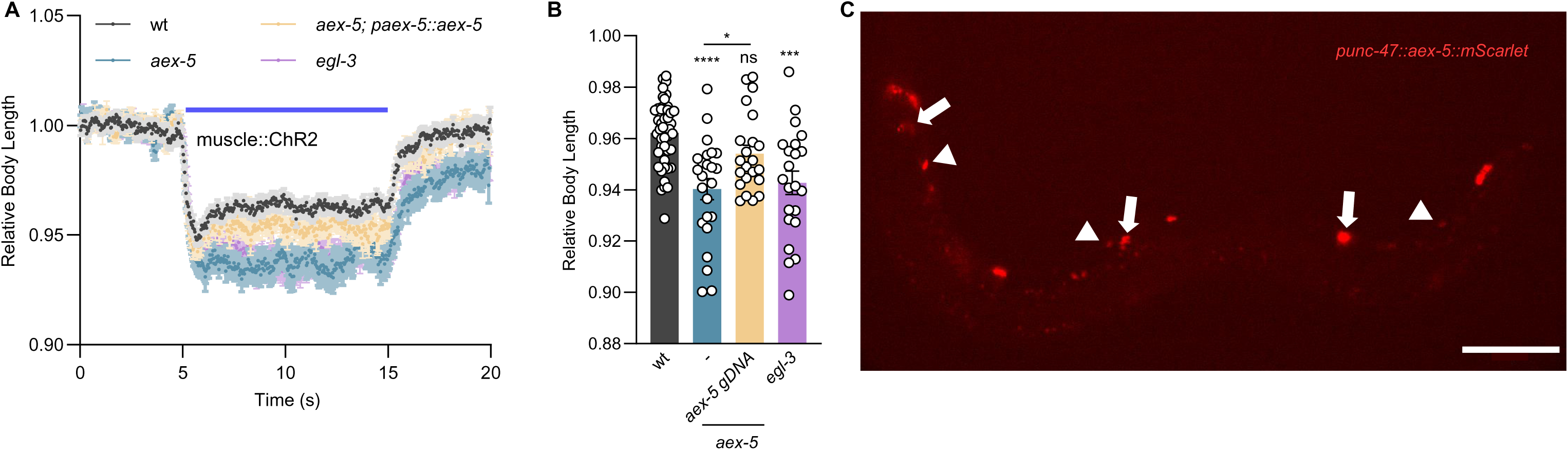
AEX-5 is released from GABAergic neurons. Related to Figure 2. **A, B)** Measurements of body length change, induced by muscular ChR2 activation using 65 µW/mm^2^ blue light stimulation. *aex-5* genomic rescue was done by specifically expressing AEX-5 under its own promoter. Animal number tested n = 46, 24, 22, 22, from left to right columns, respectively. **C)** Coelomocytes fluorescence, resulting from expression (and secretion) of mCherry-tagged AEX-5 in GABAergic neurons (using the *unc-47* promoter). White arrows indicate the coelomocyte in the middle part of the animal. White arrowheads indicate the ventral cord GABAergic neurons. Scale bar 100 µm. Data presented as mean ± SEM. Statistical significance for multiple-group datasets comparison was determined using one-way ANOVA with Tukey-correction. *, ***, and **** indicate p < 0.05, p < 0.001, and p < 0.0001, respectively.

**Figure S2.**
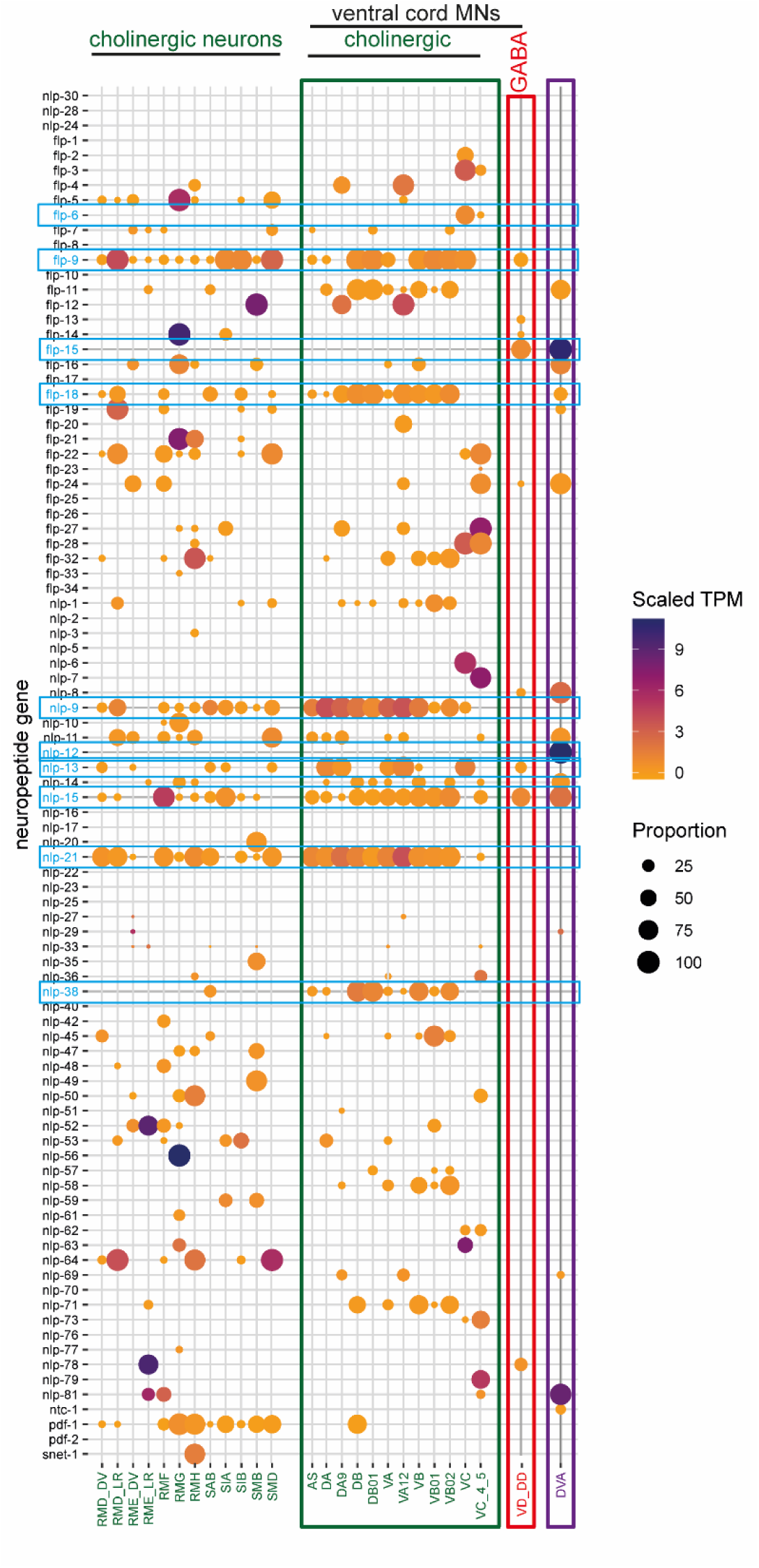
Expression data summary of neuropeptides expressed in MNs. Related to Figure 3. **A)** Heatmap representing the expression patterns and levels of neuropeptide genes in different neuron types were generated from CeNGENApp (https://cengen.shinyapps.io/CengenApp/). The initial exported plot was cropped to show cholinergic neurons, GABAergic neurons, and the cholinergic interneuron DVA. Scaled TPM (transcripts per kilobase million) and proportion data refer to all neurons of *C. elegans*, not only the subset shown.

**Figure S3.**
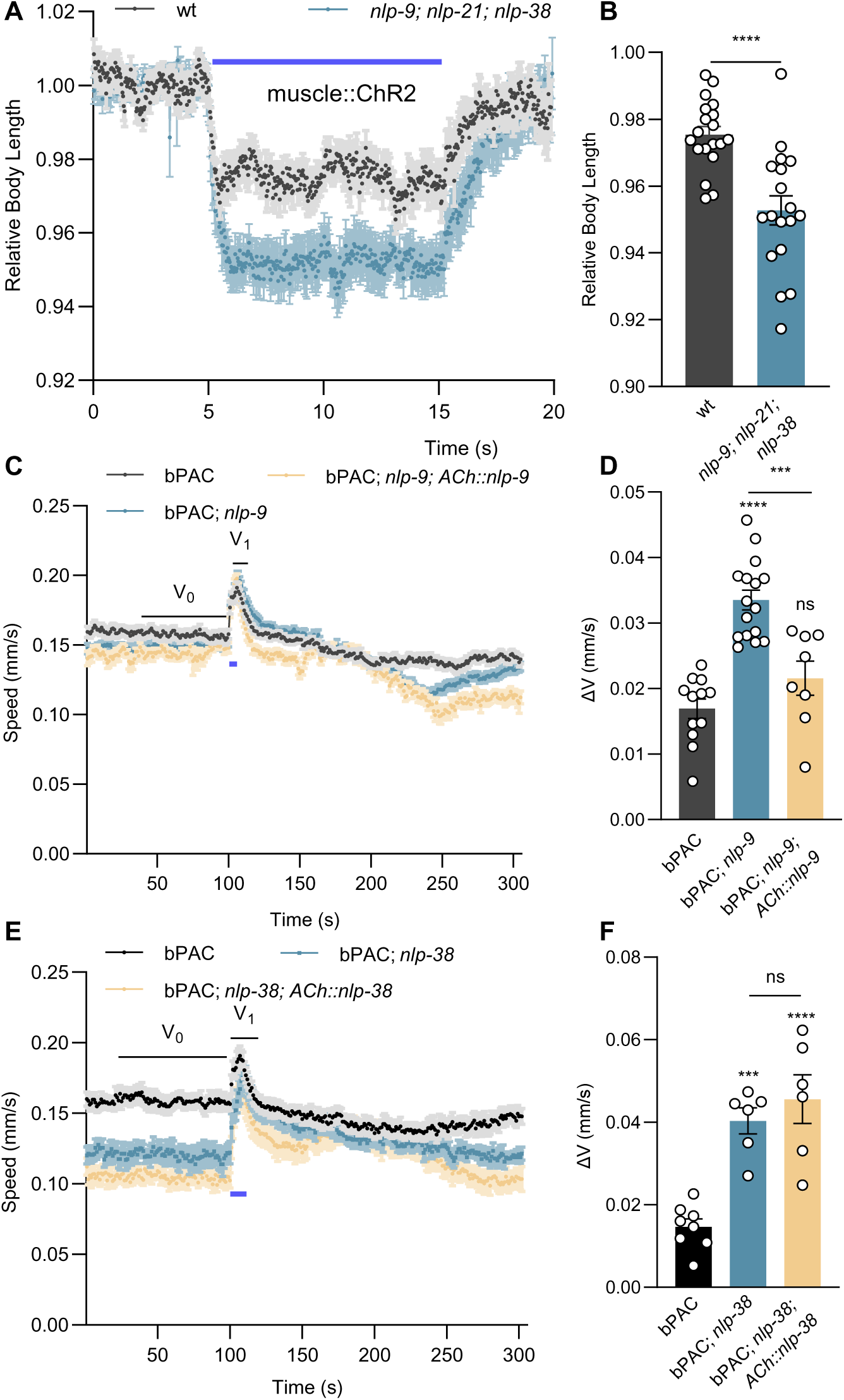
Loss of expression of NLP-9, NLP-21 and NLP-38 in cholinergic neurons increases postsynaptic excitability. Related to Figure 4. **A, B)** Measurements of body length induced by muscular ChR2 activation using 65 µW/mm^2^ blue light stimulation in wt (n = 19) and *nlp-9; nlp-21; nlp-38* triple mutants (n = 18). **C-F)** Crawling speed induced by cholinergic bPAC activation was compared in the indicated genotypes. *nlp-9* (C, D) and *nlp-38* rescue (E, F) were done by specifically expressing NLP-9 and NLP-38 from the cholinergic promoter *unc-17*. Animal number tested in (D) n = 60-80, in N = 12, 16, 8 experiments, and in (F) n = 60-80, in N = 8, 6, 6 experiments, from left to right, respectively. All data are presented as mean ± SEM. Statistical significance for two-group datasets and multiple-group datasets comparison was determined using unpaired t test and one-way ANOVA with Tukey-correction respectively. ***, and **** indicate p < 0.001, and p < 0.0001, respectively.

**Figure S4.**
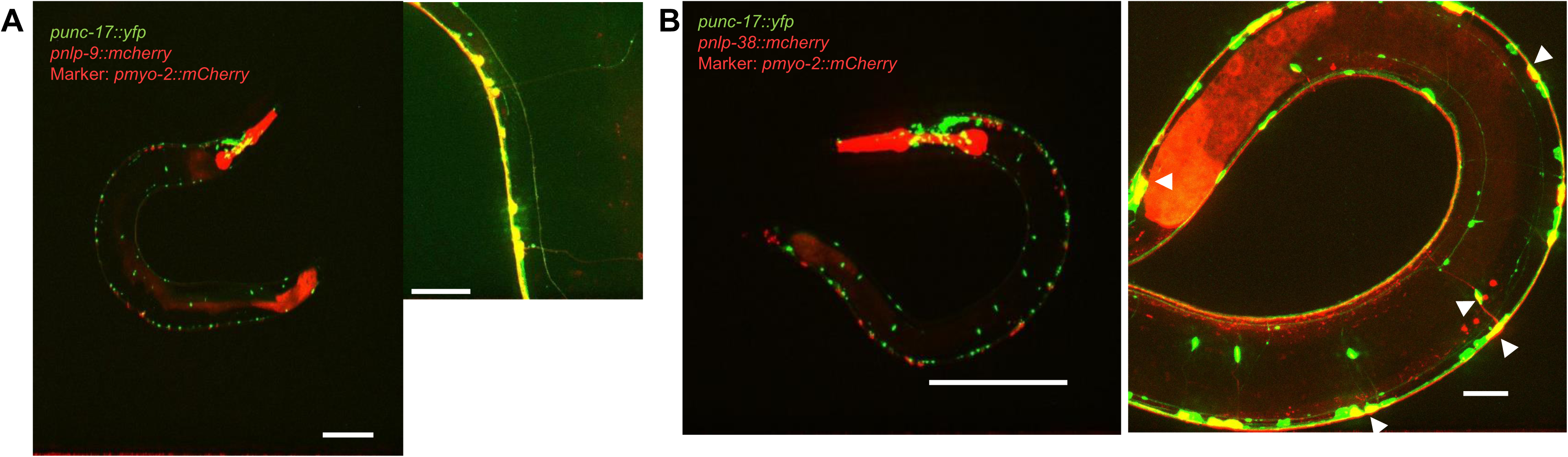
Anatomical expression patterns of *nlp-9* and *nlp-38*. Related to Figure 4. **A, B)** Representative images of *nlp-9* and *nlp-38* expression pattern indicated by mCherry fluorescence driven by the *nlp-9* and *nlp-38* promoters, respectively and their colocalization with the fluorescence of YFP, expressed in cholinergic neurons using the *unc-17* promoter. White arrowheads indicate colocalization of cell bodies of *nlp-9* or *nlp-38* expressing neurons and ventral cord cholinergic MNs. Scale bars 100 and 20 µm, respectively.

**Figure S5.**
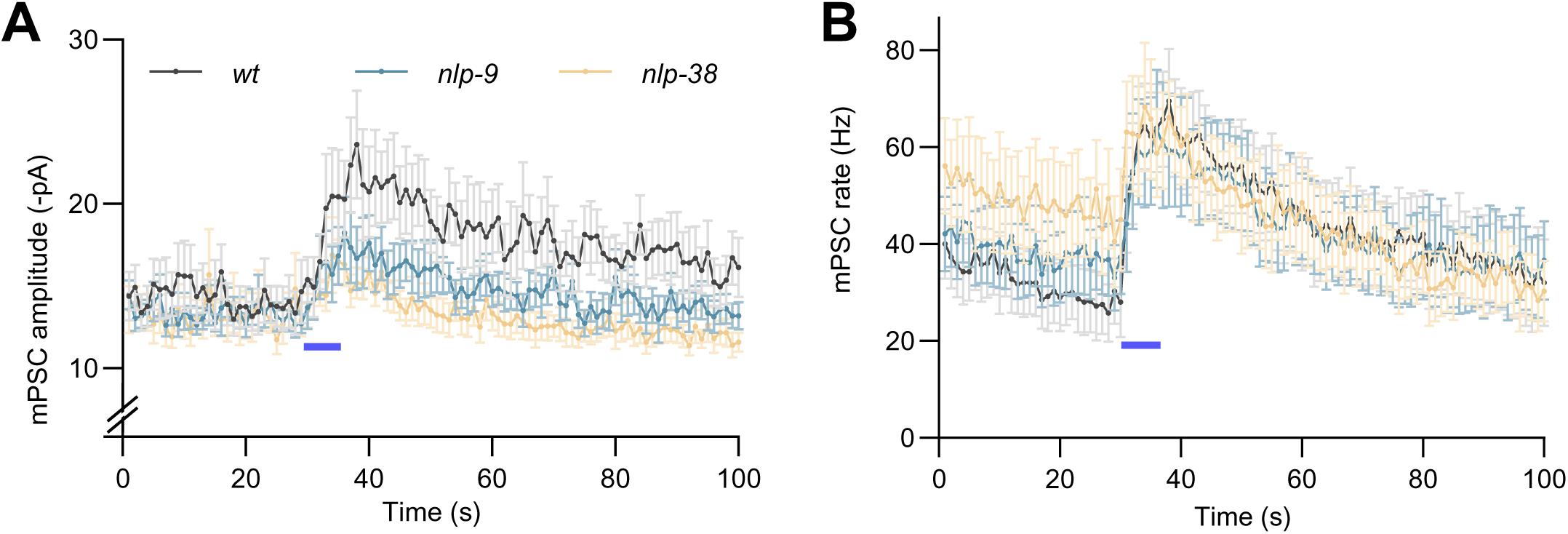
Analysis of (m)PSC amplitude and rate, induced by bPAC stimulation, in *nlp-9* and *nlp-38* mutants. Related to Figure 5. (A-B) Mean ± SEM mPSCs, recorded from dissected body wall muscle of adult worms in wt (n = 12), *nlp-9* (n=13) and *nlp-38* mutants (n = 8). Blue bar indicates stimulation of bPAC in cholinergic neurons. Data as in Fig. 5E, F, not normalized.

**Figure S6.**
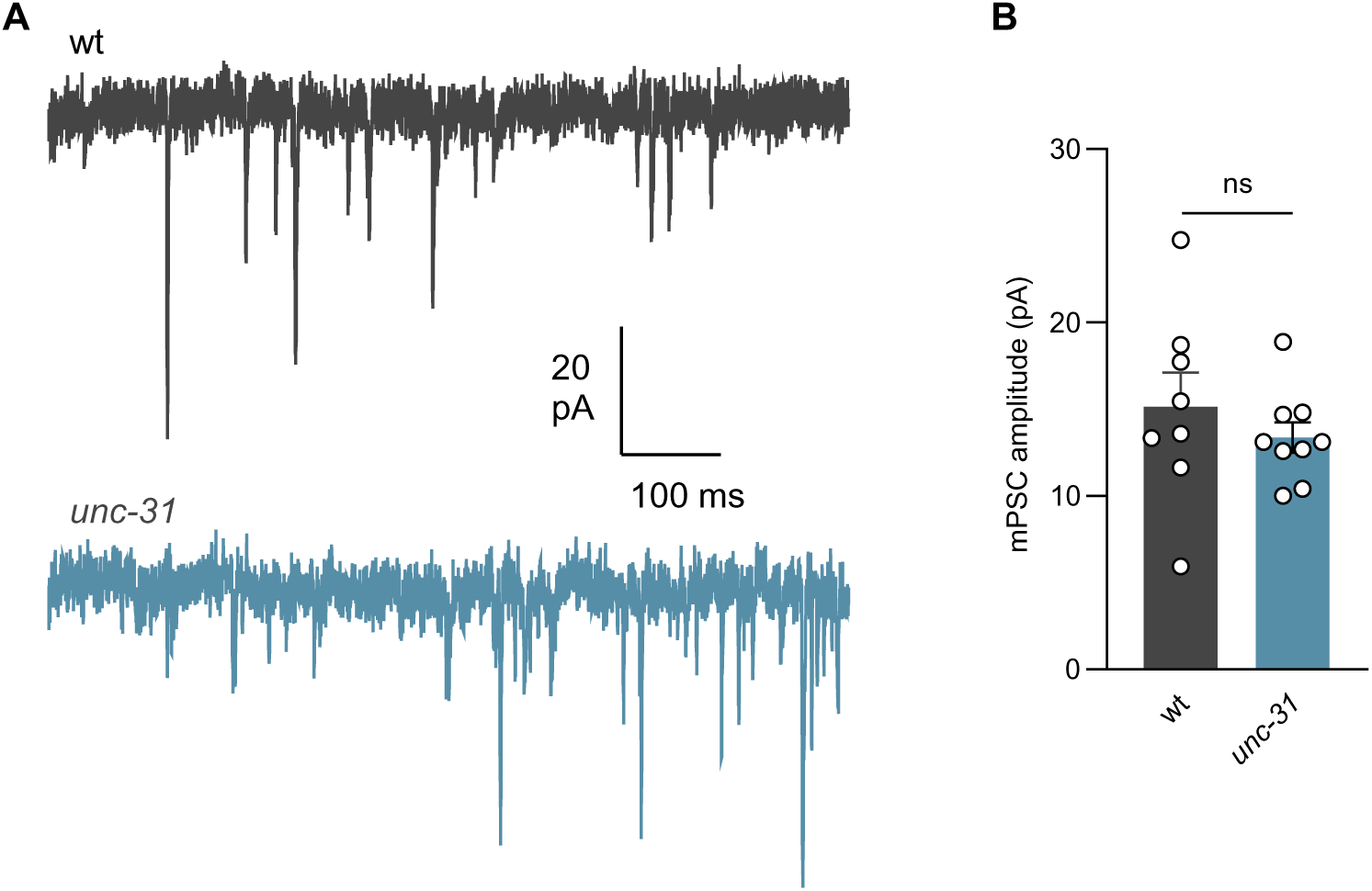
Example current traces and mPSC amplitude in wt vs. *unc-31* mutants. Related to Figure 6. **A, B)** Basal, unstimulated mPSCs recorded from dissected BWM cells of adult worms in wt (n = 8) and *unc-31* mutants (n = 9). Representative traces of mPSCs (A) and summary data of mPSC amplitudes (B) are shown. All data are presented as mean ± SEM. Statistical significance comparison was determined using unpaired t test. ns = not significant.

**Figure S7.**
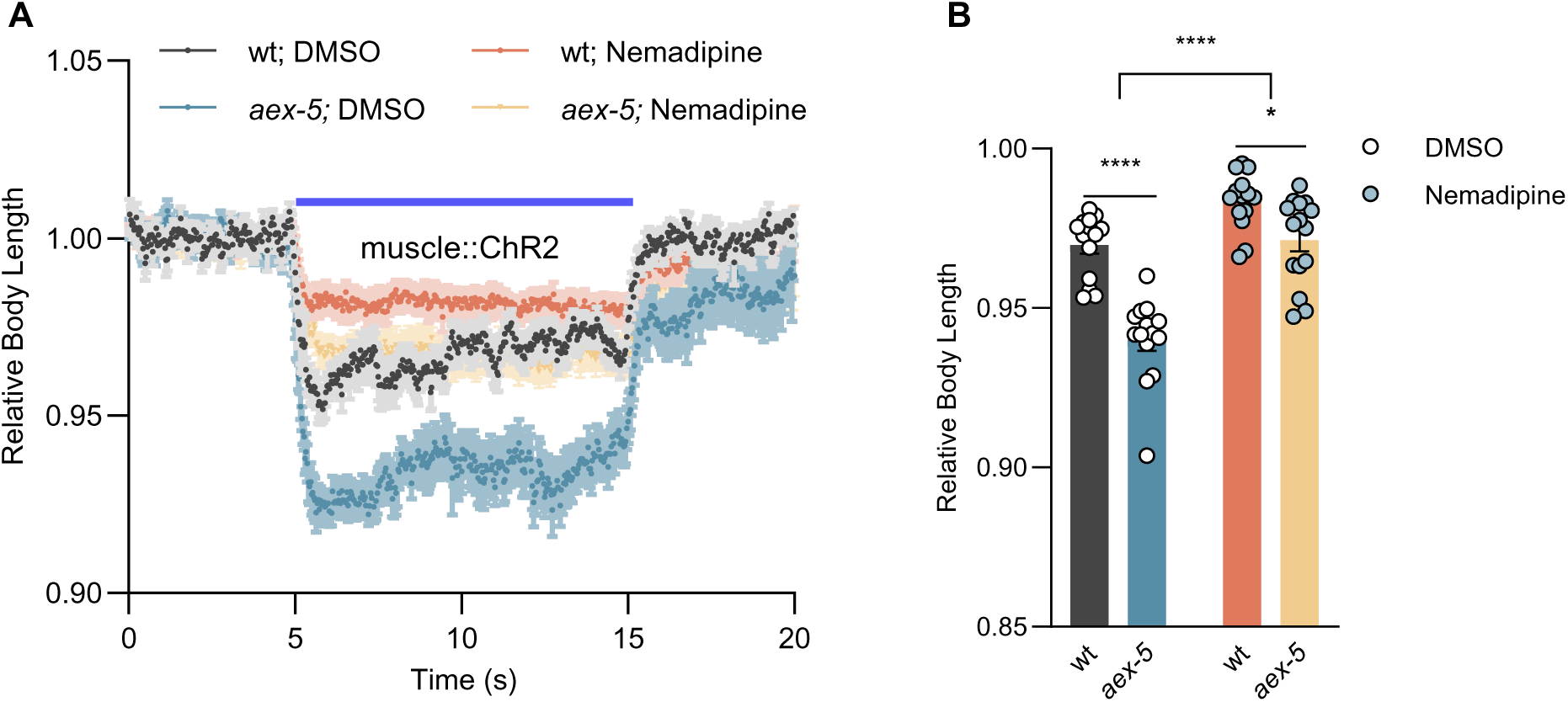
Increased muscular excitability in *aex-5* mutant is reverted by CaV1 block. Related to Figure 7. **A, B)** Body contraction induced by muscular ChR2 activation using 65 µW/mm^2^ blue light stimulation was compared in the indicated genotypes and treatments. Relative body lengths after treatment with the CaV1 specific inhibitor nemadipine are shown. Number of animals tested, n = 14, 14, 14, 15, from left to right columns, respectively. Data are presented as mean ± SEM. Statistical significance comparison was determined using one-way ANOVA with Tukey-correction and two-way ANOVA with Sidak’s multiple comparisons test. *, and **** indicate p < 0.05, and p < 0.0001, respectively.

**Figure S8.**
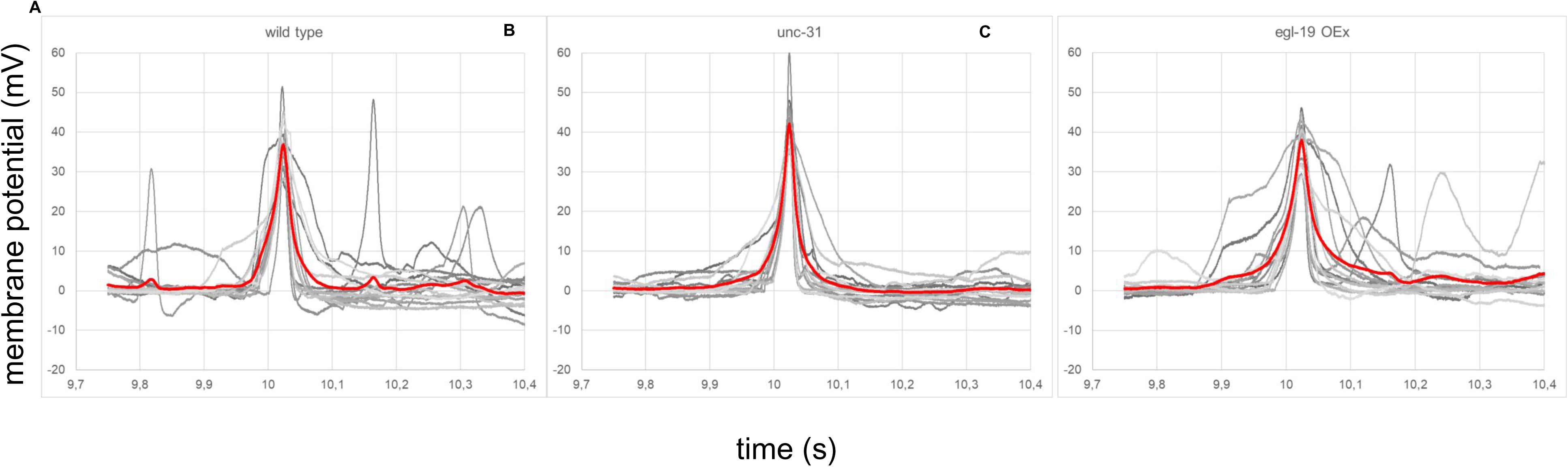
Enhanced AP amplitude in *unc-31* mutants; delayed and prolonged APs in animals overexpressing EGL-19 CaV1. Related to Figure 8. **A-C)** Individual (gray) and mean (red) traces of membrane potential changes recorded from BWM cells after 20 pA current-step induced at 10.01 s, in wt (A), *unc-31* (B), and EGL-19 over-expressing (OEx) animals (C) are shown, from n = 15, 15, and 14 animals, respectively. Note that the traces were aligned to the time of the peak amplitude, and replotted, thus they appear shifted to earlier times.

