## Supplemental Table 1 for "Loss of neuropeptidergic regulation of cholinergic transmission induces CaV1-mediated homeostatic compensation in muscle cells"

| neuropeptide | protein sequence | EGL-3 (PC2) cleavage sites | AEX-5 (PC1) cleavage sites |
|---|---|---|---|
| | | RR or KR | RXXR |
| *nlp-9a* | MDRFATRFIALLLVLLQIGSIFATPIAEAQGAPEDVDDRRELEKRGGARAFYGFYNAGNSKRDQAAALPYYLYEKRGGGRAFNHNANLFRFDKRGGGRAFAGSWSPYLERFYDYKRSSYPVYFSDNSYY | yes | no |
| *nlp-9b* | MDRFATRFIALLLVLLQIGSIFATPIAEAQGAPEDVDDRRELEKRGGARAFYGFYNAGNSKRDQAAALPYYLYEKRGGGRAFNHNANLFRFDKRGGGRAFAGSWSPYLERDNSYY | yes | no |
| *nlp-38a* | MQLIHFIVGLAMLISLSLAASDDRVLGWNKAHGLWGKRSVQEASQDKRTPQNWNKLNSLWGKRSASSFDDDYTTENGDDDVTMLYKRSPAQWQRANGLWGR | yes | no |
| *nlp-38b* | MQLIHFIVGLAMLISLSLAASDDRVLGWNKAHGLWGKRSVQEASQDKRTPQNWNKLNSLWGKRSASSFDDDYTTENGDDDVTMLYKRSNLSPRFLGRMTFARIPKISPAQWQRANGLWGR | yes | no |
| *flp-15* | MQFSTLIRVAVFAVLAIATLADYDDNSVGTIPVAVDLDYFSNYVKKGGPQGPLRFGKRRGPSGPLRFGKRSSFHVAPAAEDVASWYQ | yes | no |
| *nlp-15* | MPSSSSSSSFFAAVLLVIVMMSTVESAAVRLRPVGSLFFLNRPHEKRAFDSLAGSGFDNGFNKRAFDSLAGSGFGAFNKRAFDSLAGSGFGAFNKRAFDSLAGSGFSGFDKRAFDSLAGQGFTGFEKRAFDTVSTSGFDDFKL | yes | no |
| *nlp-21a* | MRNSLFTTLFFGLAALVMVLNAQYTSELEEDEKRGGARAMLHKRGGARAFSADVGDDYKRGGARAFYDEKRGGARAFLTEMKRGGARVFQGFEDEKRGGARAFMMDKRGGGRAFGDMMKRGGARAFVENSKRDEDWVIRPFEDDRLEKRGGGRSFPVKPGRLDD | yes | no |
| *nlp-21b* | MRNSLFTTLFFGLAALVMVLNAQYTSELEEDEKRGGARAMLHKRGGARAFSADVGDDYKRGGARAFYDEKRGGARAFLTEMKRGGARVFQGFEDEKRGGARAFMMDKRGGGRAFGDMMKRGGARAFVENSKRDEDWVIRPFEDDRLGVF | yes | no |
| *nlp-12* | MLRHHSCALLMLILVFVEVFATQSPTFDRQDRDYRPLQFGKRDGYRPLQFGKRDYRPLQFGKRSSGSSGPVVLEPIWEWQ | yes | yes |
