## Supplemental Table 2 for "Loss of neuropeptidergic regulation of cholinergic transmission induces CaV1-mediated homeostatic compensation in muscle cells"

Table S2

| Plasmid | Information |
|---|---|
| pJS10 | *punc-17::nlp-38::SL2::mCherry* |
| pJS11 | *pnlp-38::nlp-38::SL2::mCherry* |
| pJS12 | *punc-17::nlp-9::SL2::mCherry* |
| pJS13 | *pnlp-9::nlp-9::SL2::mCherry* |
| pJS19 | *pacr-2::nlp-9::mCherry* |
| pJS22 | *egl-19 RNAi* |
| pJS30 | *pmyo-3::unc-31 cDNA::SL2::mCherry* |
| pJS31 | *punc-47::unc-31 cDNA::SL2::mCherry* |
| pJS32 | *prab-3::unc-31 cDNA (KG#121)* |
| pJS36 | *punc-17::unc-31 cDNA::SL2::mCherry* |
| pJS45 | *prab-3::unc-31 cDNA::SL2::gfp* |
| pJS48 | *paex-5::aex-5 cDNA::SL2::gfp* |
| pJS52 | *punc-47::aex-5 cDNA::gfp* |
| pJS56 | *punc-47::gfp* |
| pJS54 | *pmyo-3::aex-5 cDNA::P2A::gfp* |
| pJS59 | *pset-18::egl-19b (KP#2460)* |
| pJS60 | *pmyo-3::egl-19b::P2A::gfp* |
| pJS61 | *punc-47::aex-5 cDNA::P2A::gfp* |
| pJS65 | *punc-47::nlp-9::SL2::mCherry* |
| pJS69 | *punc-17::aex-5 cDNA::P2A::gfp* |
